# Epigenetic then genetic variations underpin rapid adaptation of oyster populations (*Crassostrea gigas*) to Pacific Oyster Mortality Syndrome (POMS)

**DOI:** 10.1101/2023.03.09.531494

**Authors:** Janan Gawra, Alejandro Valdivieso, Fabrice Roux, Martin Laporte, Julien de Lorgeril, Yannick Gueguen, Mathilde Saccas, Jean-Michel Escoubas, Caroline Montagnani, Delphine Destoumieux-Garzón, Franck Lagarde, Marc A. Leroy, Philippe Haffner, Bruno Petton, Céline Cosseau, Benjamin Morga, Lionel Dégremont, Guillaume Mitta, Christoph Grunau, Jeremie Vidal-Dupiol

## Abstract

Disease emergence is accelerating in response to human activity-induced global changes. Understanding the mechanisms by which host populations can rapidly adapt to this threat will be crucial for developing future management practices. Pacific Oyster Mortality Syndrome (POMS) imposes a substantial and recurrent selective pressure on oyster populations (*Crassostrea gigas)*. Rapid adaptation to this disease may arise through both genetic and epigenetic mechanisms. In this study, we used a combination of whole exome capture of bisulfite-converted DNA, next-generation sequencing, and (epi)genome-wide association mapping, to show that natural oyster populations differentially exposed to POMS displayed signatures of selection both in their genome (single nucleotide polymorphisms) and epigenome (CG-context DNA methylation). Consistent with higher resistance to POMS, the genes targeted by genetic and epigenetic variations were mainly related to host immunity. By combining correlation analyses, DNA methylation quantitative trait loci, and variance partitioning, we revealed that a third of the observed phenotypic variation was explained by interactions between the genetic sequence and epigenetic information, ∼14% by the genetic sequence, and up to 25% by the epigenome alone. Thus, as well as genetic adaptation, epigenetic mechanisms governing immune responses contribute significantly to the rapid adaptation of hosts to emerging infectious diseases.

## Introduction

There has been a substantial increase in the emergence of non-human pathogens (epizootics) resulting from human-linked activities, including anthropogenic-driven climate change, pollution, habitat fragmentation, over-exploitation, local biodiversity impoverishment, and transport of living organisms ^1-3^. Some marine epizootics have significantly disturbed ecosystems or social-ecological systems, when affecting host species of ecological or economic interest ^4^. Understanding how host populations can adapt rapidly to emerging infectious diseases will be essential for developing effective and ecologically appropriate management practices.

Host-pathogen interactions are characterized by reciprocal selective pressures that both partners impose on each other, and emerging diseases present an opportunity to study rapid selective evolutionary processes. Recent hypotheses propose that the natural phenotypic variation induced by host-pathogen selective pressures could be driven by both genetic and non-genetic components ^5,6^. Pacific oyster mortality syndrome (POMS) represents an opportunity to better understanding rapid host adaptation to an emerging pathogen. The Pacific oyster (*Crassostrea gigas*) is the most widely farmed oyster worldwide and one of the main marine resources produced by aquaculture. However, since the emergence of the Ostreid Herpes virus 1 microvariant (OsHV-1 µVar) in 2008, juvenile oysters living in high densities such as in farming area face an annual rate of mortality ranging from 40–100% worldwide ^7,8^.

POMS is a polymicrobial disease, initiated by OsHV-1 µVar virus infection, which induces lethal bacteremia ^9^. Temperature ^10^ and food availability ^11^ facilitate the establishment and/or development of POMS, by altering oyster physiology. Disease susceptibility has a heritable component^12-14^. Genome-wide association studies (GWA) have revealed that disease resistance has a polygenic architecture^14-16^. But oyster resistance to POMS is also dependent of oyster life-history, such as age ^17,18^, and past exposure to pathogen elicitors ^19,20^. Exposure to non-pathogenic microbes ^21^ also had an effect, with microbial exposure being associated with epigenetic modifications (i.e. DNA methylation) transgenerationally transmitted to offspring ^21^. These results suggest that natural oyster populations experiencing POMS constitute a useful host-pathogen system for studying the genetic and epigenetic mechanisms underlying rapid adaptation ^22^.

In this study, we have started to identify the genetic and epigenetic signatures of POMS exposure and their relative contributions to the ongoing adaptation to POMS. To do so, we used a whole exome capture approach to jointly study one component of the genetic variation (single nucleotide polymorphism, SNP) and one component of the epigenetic variation (DNA methylation in the CG context, CpG). The genetic and epigenetic underpinning of natural variation in oyster resistance to POMS were detected by carrying out GWA and epigenome-wide association (EWA) mapping, respectively. Subsequent correlation analyses, methylation quantitative trait loci (MethQTL) mapping and variance partition methods (Redundancy Analysis, RDA) allowed us to quantify the relative contribution of genetic and epigenetic variations underlying adaptation to POMS.

## Results

### Differing POMS resistance in wild oyster populations

To phenotype POMS resistance, four wild oyster populations from a “non-farming area” (B1-B4) and two from a “farming area” (B5 and B6) in the *Rade de Brest*, north-west France, were sampled (**Fig. 1A** and **Table S1**). The collected oysters were acclimatized in experimental tanks and were subjected to experimental infection (randomized complete block design) by cohabitation with infected donor oysters from two dedicated oyster families (H12, NSI) originating from a IFREMER hatchery (**Fig. 1B** and **Data S1**). A lower risk of mortality was detected for the farming area (B5 and B6) oyster populations, compared to the four populations from non-farming area (B1-B4) population (Log-rank test *P* < 0.001, **Table S2**). Resistance to disease reached > 94.9% in the farming area populations (**Fig. 1C**), whereas susceptibility was high (71.5–49.1%) in the non-farming area (**Fig. 1C**). This phenotype was coded either as a binary trait with 0 and 1 corresponding to susceptible and resistant individuals, respectively, or as a semi-quantitative trait corresponding to the survival time (expressed in hours) of an individual after its exposure to the OsHV-1 µVar (i.e. the whole duration of the experiment for the resistant oysters) (**Data S1**).

**Figure 1.**
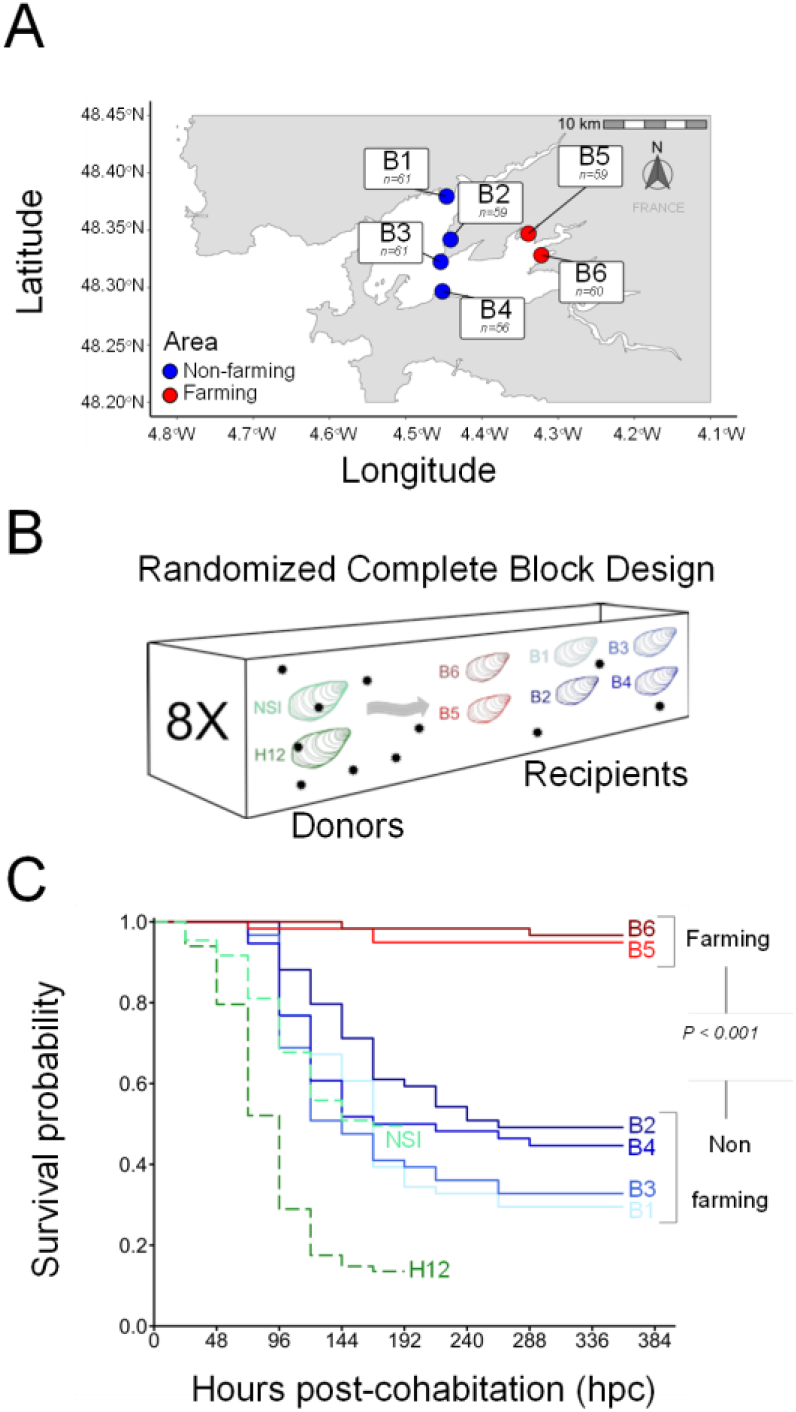
Experimental design and survival in recipients oysters after to the exposure of Pacific Oyster Mortality Syndrome (POMS). A) Oyster populations sampled in four non-farming (low density, > 20 individuals/m^2^ and no POMS reported) and two farming areas (high density with typical oyster bed of hundreds individuals/m^2^ and annual POMS reported) in the Rade de Brest (France). The n represents the number of oysters sampled in each location (Total size experiment N=356). B) Experimental infection by cohabitation was performed using a randomized complete block design (eight tanks). Donor oysters (NSI and F12) were injected with 200 µl of OsHV-1 µVar suspension and placed with the recipient oysters (B1 to B6) to induce natural disease transmission for evaluating susceptibility to POMS in the recipient oysters. C) The Kaplan– Meier survival curves of the donor (dashed line) and recipient oysters (solid line) during the cohabitation experiment. Note that donor oysters were removed from the tanks at 196 hours post-cohabitation (hpc).

Mortalities among donor oysters (H12, NSI) started 24 hours post-cohabitation (hpc). At 192 hpc, the survival rate dropped to 13.5% and 49.5% for H12 and NSI donors, respectively (**Fig. 1C**). Quantification of OsHV-1 μVar in the experimental tanks showed that viral excretion reached 1,755 ± 429 genome copies/µL seawater (mean ± SD) at 24 hpc and peaked at 72 hpc (7,185 ± 1,856 genome copies/µL). No significant difference in the load of excreted virus was observed between the experimental tanks (Kruskal-Wallis, *P* = 0.24; **Fig. S1**). Mortalities in recipient oysters (oyster from B1-B6 populations) commenced at 72 hpc (**Fig. 1C**) and did not differ significantly among the eight replicate tanks (Log-rank test *P* = 0.61) (**Fig. S2**). Thus, the farming area oyster populations, which have been subjected to high pathogen pressure, have become resistant to POMS disease, whereas the non-farming area populations, which have not experienced the disease, were highly susceptible ^23^.

### POMS resistance is associated with immune pathways

To characterize genetic (SNP) and epigenetic (CpG DNA methylation) variation in both susceptible and resistant oysters, we performed an exome-capture experiment using bisulfite-converted DNA. The exomes of 116 susceptible and 130 resistant oysters were captured and sequenced. On average, sequencing resulted in 0.5–60 million paired end (PE) reads per sample (mean ± SD = 26 ± 1 millions), with the six samples that displayed less than 7.8 millions PE reads being discarded. On average, 59.3 ± 2.7% of reads were uniquely mapped to the *C. gigas* genome and the bisulfite conversion efficiency was 99.6 ± 0.1% (**Data S2**). SNPs and DNA methylation-calling resulted in 5,110,093 SNPs and 3,449,600 CpGs for the 240 oyster analysed. After applying filtering criteria for GWA and EWA mapping, data from 102 susceptible and 118 resistant oysters, characterized by 214,263 SNPs and 635,201 CpGs, were used in subsequent analysis (**Data S2**). No signatures of population structure were detected at the genomic or epigenomic levels (**Fig. S3**). A small percentage of the genetic (R^2^ = 2.3%, *P* = 0.091) and epigenetic variance (R^2^ = 2.4%, *P* < 0.001) was explained by differences among the six populations using (PERMANOVA).

GWA mapping revealed one SNP significantly associated with the binary trait: scaffold1832_479264; A>T; *P* = 5.53E^-08^ (**Fig. 2A, Fig. S4A**, and **Data S3**) and one SNP with the semi-quantitative trait: scaffold364_478394; C>T; *P* = 1.13E^-07^ (**Fig. 2B, Fig. S4B**, and **Data S4**). The SNP associated with the binary trait was mapped on chromosome 6 in a gene encoding the SUMO-activating enzyme subunit 2 (CGI_10018487), and the SNP associated with the semi-quantitative trait was mapped on the chromosome 4 in a gene of unknown function (CGI_10022698). Given the low number of significant SNPs identified and the polygenic architecture of POMS resistance ^14-16^, we extended our analysis to suggestive SNPs (*P*-value < 0.0005), which led to the identification of 113 and 112 SNPs associated with the binary and semi-quantitative traits, respectively **(Data S3** and **S4)**. Among these 225 SNPs, 186 of which are non-redundant, 39 were in common, and 74 and 73 were specific to the binary and semi-quantitative traits, respectively (**Fig. 2C** and **Data S5**). The 186 SNPs were located in 155 genes, with 37 genes in common, and 58 and 60 being specific to the binary and semi-quantitative traits, respectively (**Fig. 2C** and **Data S6**).

**Figure 2.**
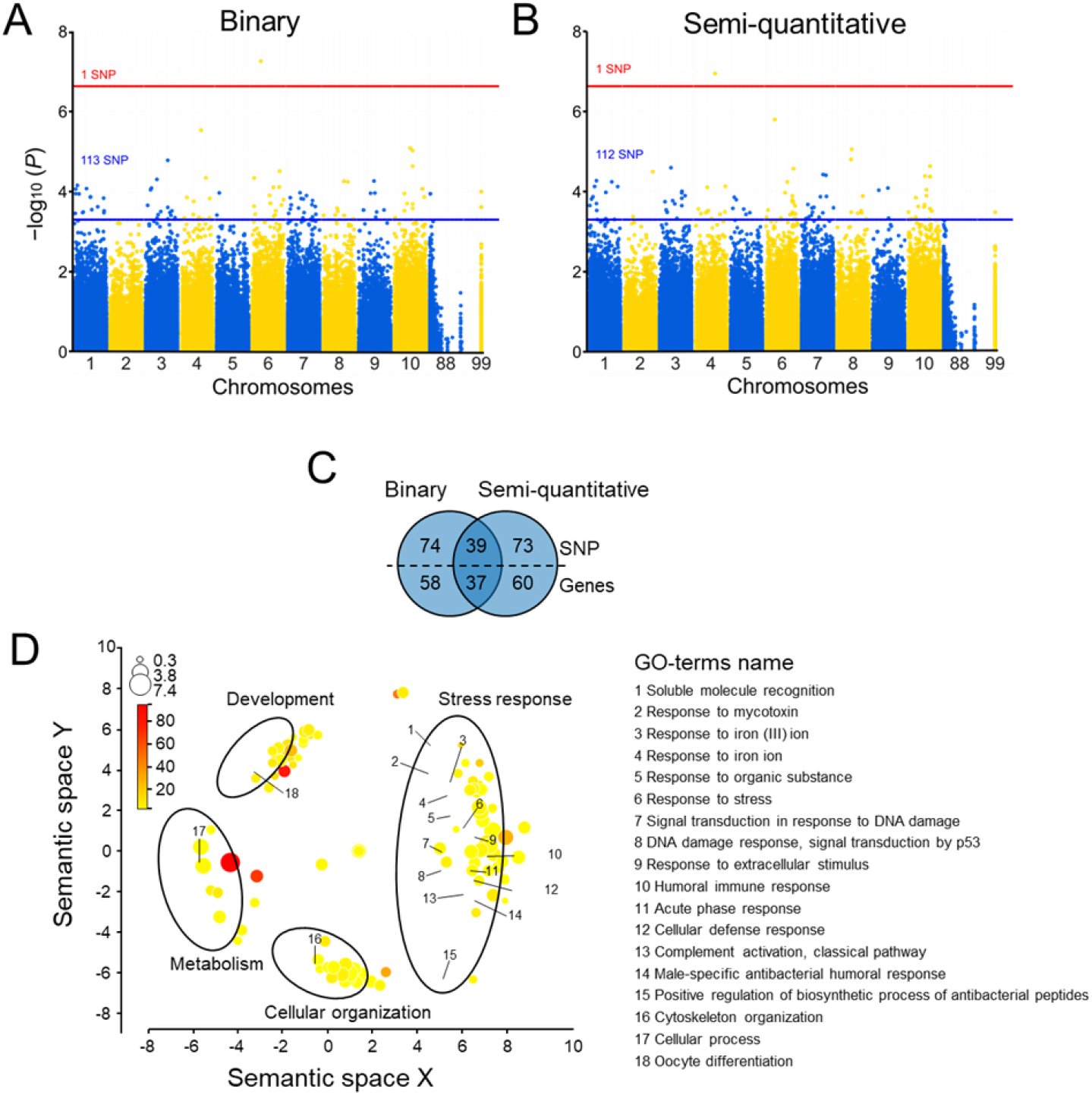
Genetic variation associated (SNPs) with resistant phenotype to Pacific Oyster Mortality Syndrome (POMS). Result of GWA mapping with A) binary phenotype or B) semi-quantitative trait of POMS resistance. Chromosomes 88 and 99 correspond to unknown regions in the genome. The red line represents the threshold for a False Discovery Rate < 0.05 (significant SNPs), and the blue line the threshold for a *P* value < 0.0005 (suggestive SNPs). C) Venn diagram comparing the number of SNPs and genes between associations obtained from the binary and semi-quantitative phenotype. D) Gene enrichment analysis (GO-terms) of the Biological Process (BP) from the set of genes displaying significant and suggestive SNPs for the binary and semi-quantitative GWA mapping. GO-terms are distributed in multidimensional semantic similarities. The size of the circles (log10 size) and the color saturation (log10 Fisher’s *P* value) indicate the number of genes represented and the significance value for each GO term, respectively.

Enrichment analysis of Gene Ontology (GO-terms) revealed the enrichment of four groups of Biological Process (BP) category: development, metabolism, cellular organization, and stress response (**Fig. 2D** and **Data S7**). In the last category, key stress response BPs were retrieved, including: Cellular defence response, humoral immune response, complement activation classical pathway, positive regulation of biosynthetic process of antibacterial peptides, response to extracellular stimulus, signal transduction in response to DNA damage, and DNA damage response (**Fig. 2D**). In these BPs including immune processes, we identified genes involved in the JAK/STAT pathway (e.g. PRMT5, AIMP1, UBA2, and DCST1), the STING/RLRs pathway (e.g. TRIM33, TRAF3), the TLR/NF-KB pathway (e.g. MIB2, MyD88, PRGP, TBK1), the RNAi pathway (Dicer), and pathogen recognition (e.g. C1q, DSCAM, MR) (**Fig. 3A**). Thus, POMS events have selected genetic variation in multiple genes of key immune pathways.

**Figure 3.**
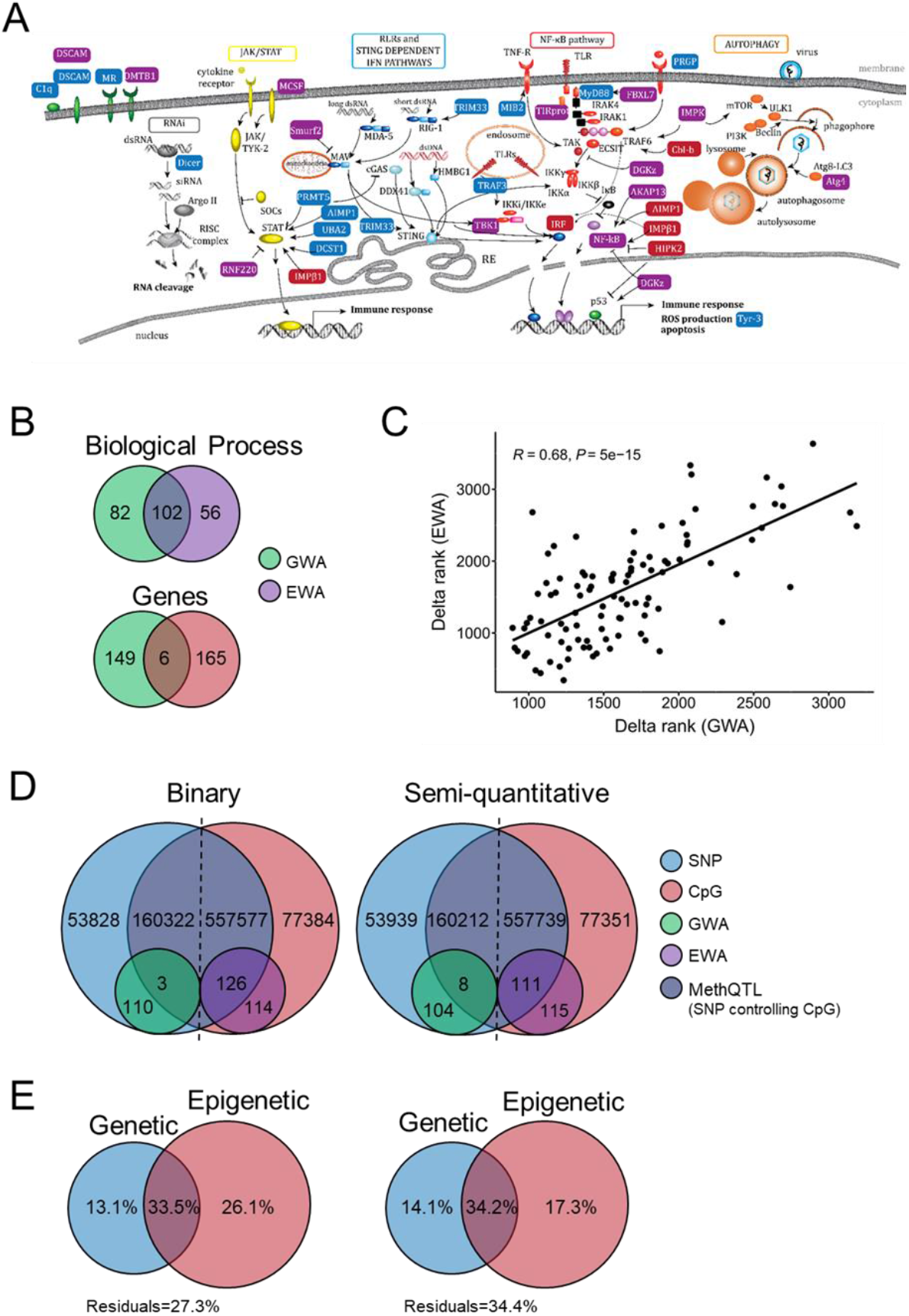
Genes involved in innate immune pathways. A) Genes involved in innate immune signalling pathways identified with genetic variation only (SNP, blue, 14 genes), epigenetic variation only (CpG, red, 4 genes), and genetic and epigenetic variation (violet: SNP+CpG 1 gene; methQTL 12 genes). B) Venn diagram comparing the GO-terms of Biological Process and genes between GWA and EWA mapping. C) Correlation between the delta rank of the GO-terms significantly enriched in the genes set obtained with GWA and EWA analysis. D) Venn diagrams comparing the results of MethQTL mapping obtained with the binary (left) or semi-quantitative (right) phenotype as covariate. E) Results of Redundancy Analysis (RDA) performed to disentangle and weight the portion of phenotypic variation (binary and semi-quantitative approaches) explained by the genetic variation, the epigenetic variation and their interaction.

To detail the epigenetic differences between susceptible and resistant oysters, we analysed the EWA mapping, which revealed that 240 (**Data S8**) and 226 (**Data S9**) differentially methylated CpGs were significantly associated with the binary and semi-quantitative traits, respectively (**Fig. 4A, 4B, Fig. S4C** and **S4D**). Among these 446 CpGs (305 being non-redundant), 161 were in common, and 79 and 65 CpGs were specific to the binary and the semi-quantitative traits, respectively (**Fig. 4C** and **Data S10**). Among the 305 non-redundant CpGs, 23 and 282 were hypermethylated and hypomethylated, respectively in resistant compared to the susceptible oysters. At the gene level, 171 genes displayed at least one differentially methylated CpG, 99 were in common, and 41 and 31 were specific to the binary or semi-quantitative traits, respectively (**Fig. 4C** and **Data S11**).

**Figure 4.**
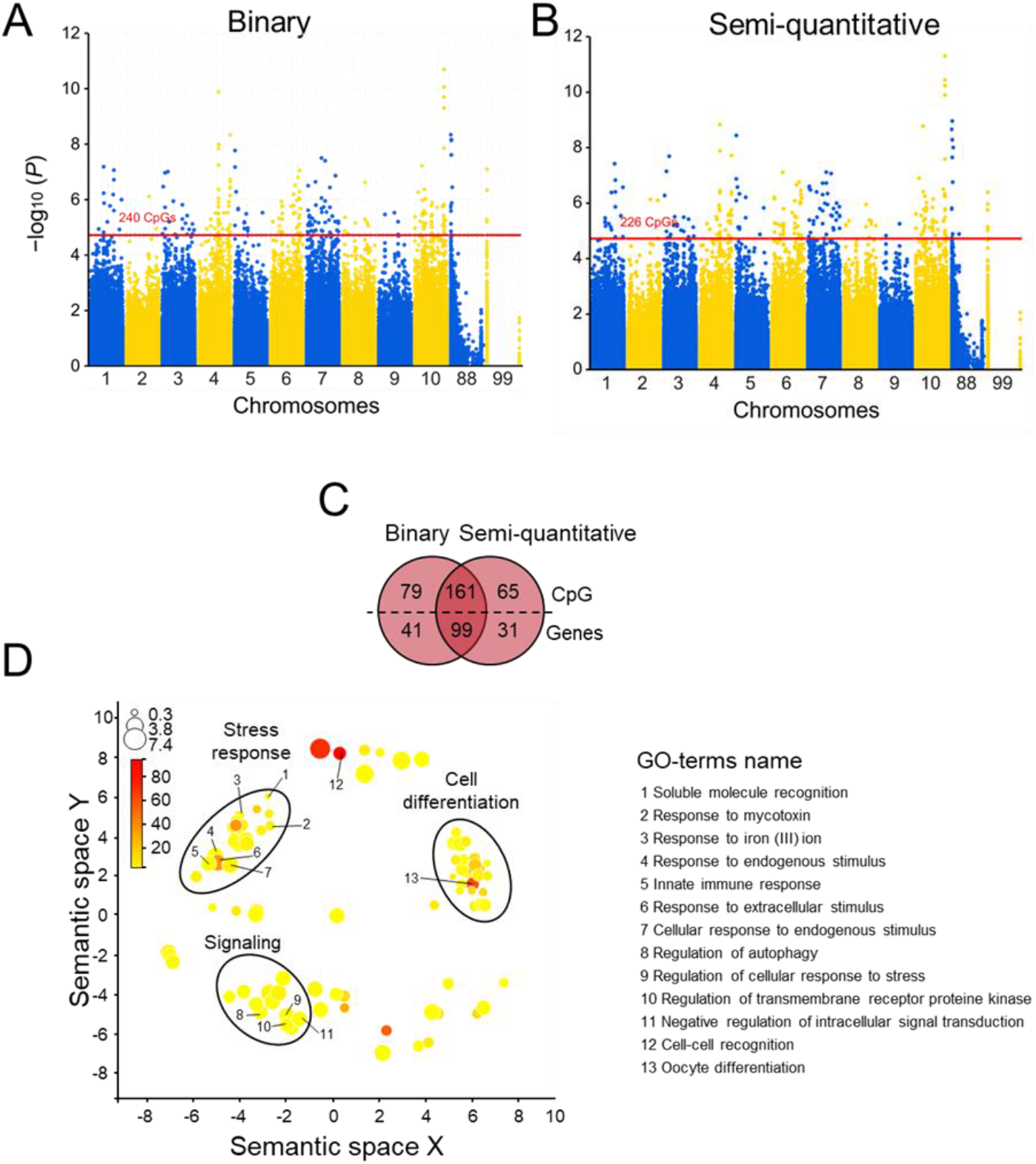
Epigenetic component associated (CpG) with the resistant phenotype to Pacific Oyster Mortality Syndrome (POMS) Result of EWA mapping with A) binary phenotype or B) semi-quantitative trait of POMS resistance. Chromosomes 88 and 99 correspond to unknown regions in the genome. The red line represents the threshold for a False Discovery Rate < 0.05 (significant CpGs) C) Venn diagram comparing the number of CpGs and genes between associations obtained from the binary and semi-quantitative phenotype. D) Gene enrichment analysis (GO-terms) of the biological process from the set of genes displaying significant and suggestive CpGs for the binary and semi-quantitative EWA mapping. GO-terms are distributed in multidimensional semantic similarities. The size of the circles (log10 size) and the colour saturation (log10 Fisher’s *P* value) indicate the number of genes represented and the significance value in each GO term, respectively.

Enrichment analysis of GO-terms revealed three main groups of BP category: cell differentiation, stress response, and signalling (**Fig. 4D** and **Data S12**). Key immune BPs were retrieved in the stress response and signalling categories: Innate immune response, response to endogenous stimulus, response to extracellular stimulus, cellular response to endogenous stimulus, regulation of autophagy, regulation of cellular response to stress, regulation of transmembrane receptor protein kinase, and negative regulation of intracellular signal transduction (**Fig. 4D**). We identified several immune genes known to be involved in JAK/STAT pathway (e.g. MCSF, RNF220, IMPβ1), the STING/RLRs pathway (e.g. Smurt2 and TBK1), the TLR/NF-KB pathway (e.g. TIRprot, FBXL7, IMPK, Cb1-b, DGKz, AKAP13, AIMP1, IMPβ1, HIPK2, TBK1, NF-kB, IRF), pathogen recognition (e.g. DMTB1) and the autophagy pathway (IMPK, ATG4) (**Fig. 3A**), similar to GWA mapping. Thus and similarly to the genetic variation, POMS events have selected epigenetic variation in multiple genes belonging to key immune pathways.

#### POMS resistance is associated to genetic and epigenetic variations in common processes

Among the 240 GO-terms enriched in the GWA and EWA mapping, 102 were in common, and 82 and 56 were specific to genetic and epigenetic variation, respectively (**Fig. 3B** and **Data S13**). The delta-rank of the 102 common GO-terms enriched in the GWA and EWA mapping showed a significant and positive correlation (Pearson correlation coefficient *R*=0.68, *P* < 0.01) (**Fig. 3C**). Of the 320 genes identified in the GWA and EWA mapping, each had at least one significant/suggestive SNP or one significant CpG (**Data S14**), and 149 were specific to genetic variation and 165 to epigenetic variation, with six genes displaying both one significant/suggestive SNP and one significant CpG (**Fig. 3B**), including TBK1, a major activator of antiviral pathways (**Fig. 3B**). Thus, resistance to POMS may select both genetic and epigenetic variations in different genes that nonetheless are involved in similar biological functions, in particular, in different genes involved in innate immunity.

#### Independent genetic and epigenetic variation

To quantify the relative contribution of genetic and epigenetic variation to the phenotypic variation we observe in POMS resistance, we tested the relationship between the matrix of pairwise genetic and epigenetic distances and detected a significant but weak correlation (Mantel statistic *rho* = 0.089, *P* = 0.0184). We also used MethQTL mapping, which identified 5,151,194 and 5,152,611 significant SNP-CpG pairs (FDR<0.05) when using the binary and the semi-quantitative trait as a covariate, respectively. Removing redundant SNP-CpG pairs, we identified 160,325 (binary) and 160,220 (semi-quantitative) SNPs associated with the DNA methylation rate of 557,703 (binary trait) and 557,850 (semi-quantitative) CpGs (**Fig. 3D** and **Table S3**), showing that a large portion (∼88%) of the epigenetic variation is under a genetic control. However, when we specifically look for the CpGs significantly associated with POMS resistance (EWA), only 126 (binary) and 111 (semi-quantitative) CpGs were involved in a methQTL, thereby suggesting that ∼50% of the significant CpGs displayed a DNA methylation rate independent of the DNA sequence (**Fig. 3D, Table S3**)

Variance partition analysis (RDA) showed that genetic and epigenetic variation jointly explained the highest percentage of phenotypic variation, with 33.5% and 34.2% for the binary and semi-quantitative traits (**Fig. 3E**). Overall, epigenetic variation explained a higher proportion of phenotypic variation (binary trait = 26.1% and semi-quantitative trait = 17.3%) than genetic variation (binary trait = 13.1% and semi-quantitative trait = 14.1% (**Fig. 3E**). Thus, most of the genetic and epigenetic variation we find associated with POMS resistance is correlated with each other. However, a fraction of the genetic and epigenetic variation remains independent, indicating that changes to the epigenome, independent of genetic changes, must underlie the selection for POMS resistance.

## Discussion

Here, we show that populations of wild oysters exposed to POMS displayed signatures of selections both in their genome (SNPs) and their epigenome (CpG DNA methylation). This selection, on genetic and epigenetic variation, targeted the same biological processes (e.g. immunity) but acted through different genes. Correlation analysis between genetic and epigenetic variation and MethQTL mapping showed that genetic and epigenetic variations are partially correlated and that genetic variation influences a large proportion of the epigenetic variation (88%). However, ∼50% of CpGs significantly associated with oyster resistance are among the 12% of epigenetic variation not controlled by genetic variation, indicating that epigenetic variation occurs independently of genetic variation. These results were confirmed by RDA analysis, showing that the expressed phenotypic variance was mainly explained (33.5-34.2%) by the interaction between genetic and epigenetic variation, a smaller fraction (17.3-26.1%) by epigenetic variation alone, and the smallest fraction (13.1-14.1%) by genetic variation alone. These results show that a host population facing recurrent and strong pathogen selection pressure experiences both genetic and epigenetic variation during rapid adaption.

In our study, we took advantage of the differential environmental selective pressure exerted by POMS on wild oyster populations, in farmed areas *vs*. non-farmed areas, to quantify the interaction and relative effect of genetic and epigenetic variations on phenotypic variation (i.e. resistant *vs*. susceptible). GWA and EWA mapping of individual oysters displaying contrasted phenotypes, enabled us to identify signatures of selection both in the genome and the epigenome. We quantified the relative effect of the genetic and epigenetic variation using genetic/epigenetic matrix correlation tests (partial Mantel test), MethQTL mapping (control of CpG methylation level by SNPs), and variance partition analysis by RDA. These analyses demonstrated that most genetic and epigenetic variation are correlated and, therefore, causally linked, given methylation of ∼88% of CpGs was significantly associated with one or more SNP. However, the 12% of CpGs that were independent of genetic control included ∼50% of the signature of selection identified by the EWA mapping. Thus, the adaptive phenotype observed in response to POMS selection involves both genetic and epigenetic variation, much of which is correlated, with genetic variation controlling much of the epigenetic variation. However, there is also independent adaptive genetic and epigenetic variation.

Significant associations between genetic or epigenetic variation and environmental parameters (e.g. temperature or salinity) have previously been reported in various invertebrates ^24-28^, with significant correlation between genetic and epigenetic variation found in six of 14 studies reviewed ^29^. Analyses of genetic control of epigenetic variation using methQTL have found that the fraction of epigenetic variation under direct control of DNA sequence variation ^25,27^,30 is highly variable, ranging from 2% in the threespine stickleback (*Gasterosteus aculeatus*) ^30^, 3% in the Olympia oyster (*Ostrea lurida*) ^27^, 19% in *Ciona intestinalis* ^25^, 70% in human (*Homo sapiens*) ^31,32^, and 88% in this study. Complementary approaches in invertebrates and vertebrates have shown that 27% of inter-individual epigenetic variation is genotype-dependent ^27^ and 24–35% of epigenetic variation is explained by additive genetic components ^30^. Our results are consistent with these studies, which, altogether, demonstrate that genetic and epigenetic variation are causally linked but also that a significant proportion of the epigenetic variation is independent of the genetic variation, and *vice versa*, such that each on its own can contribute to an adaptive phenotype.

Variance partition analysis (RDA) further supported this conclusion, showing that 33.5–34.2% of the observed phenotypic variation was explained by the interaction between the DNA sequence and epigenetic information, 26.1–17.3% by the epigenome alone, and 14.1–13.1% by the DNA sequence alone. Similar approaches had been used before to analyse gene expression variation between sister species of whitefish: Lake whitefish (*Coregonus clupeaformis)* and the European whitefish (*C. lavaretus*), both comprising sympatric benthic and limnetic specialists ^33^. There, 46.7% of gene expression variation was explained by the interaction between genetic and epigenetic variation, 4.1% by genetic variation alone, and 2.1% by epigenetic variation alone. The large differences in the relative involvement of genetic and epigenetic variation in phenotypic variation may reflect differences in mechanism acting at the macroevolutionary *vs*. microevolutionary scale. In our study, the selected oysters faced a strong environmental constraint due to a recently emerged infectious disease (14 years ago), while the whitefish study focused on an evolutionary step that occurred during the Pleistocene (∼12,000 years ago) ^33^. At the microevolutionary scale, to survive, a population under strong selection pressure will likely need to produce new phenotypes rapidly. Epigenetic variation can occur quickly, and is reversible, and therefore could allow rapid phenotype sampling, whereas genetic variation is much slower, and is almost always not reversible. The initial successful phenotypes driven by epigenetic variation could then be selected for genetic assimilation, the latter being incidentally promoted by epigenetically facilitated mutational assimilation^27,34-36^.

In our study, the non-genetic epigenetic variation associated with resistance to POMS may have arisen from environmental exposure or from random epimutations subsequently selected for by POMS. POMS interactions with the oyster immune system have demonstrated that oyster resistance/susceptibility can be influenced by both intrinsic and extrinsic factors ^37^. For example, exposure to a non-pathogenic microbiota during early life ^21^ or exposure to viral mimics (Polyinosinic: polycytidylic acid, Poly(I:C)) at the juvenile stage ^20,38^ can increase long-term immune competence, both within and across generations. Such environmental exposure induced a significant increase in resistance to POMS. Exposure to microbiota modified DNA methylation patterns ^21^, some of which were transmitted to the subsequent generation, although this second generation was never exposed to the microbiota ^21^. Changes in the epigenome induced by the environment in oysters and inheritance of DNA methylation patterns have also demonstrated in two other studies^27,28^. Thus, the environment may induce heritable epigenetic modifications in oysters which may explain long-term adaptive phenotypic traits^21,27^,28. We hypothesize that, in our study, natural microbial exposure experienced in the field may have facilitated development of resistance to POMS in the oyster population sampled in farming areas.

Exome capture, sequencing and downstream analysis performed in this study showed that the signatures of oyster resistance to POMS was carried by both genetic and epigenetic marks: 186 suggestive SNPs and 305 significant CpGs, respectively. The gene ontology enrichment analysis showed a strong enrichment of biological processes linked to immunity (**Fig. 2D** and **Fig. 4D**). Genes involved in these immune processes have been identified (**Fig. 3A**) and participate in the JAK/STAT pathways (UBA2, RNF220), the RLR/STING pathway (TRIM33, TBK1, IRF), the NF-KB pathway (TIRprot, NF-KB, MyD88), RNAi (DICER), and autophagy (ATG4), as well as several pathogen recognition receptors, such as DSCAM, Mannose Receptors, C1q, and PRGP ^39^, consistent with previous studies demonstrating the polygenic nature of POMS resistance at the genetic level ^14-16,40^. The POMS resistance phenotype involves antiviral genes and pathways, either constitutively expressed ^41^ and up-regulated faster in response to POMS in resistant families ^9^, or environmentally induced ^19-21^. These studies demonstrate that many immune system-related genes are involved in the POMS resistance phenotype, but few are common across the different oyster families or populations studied. This polygenic feature may arise from the need to produce new phenotypes rapidly and can explain why the signature of genetic and epigenetic selection is spread across a number of genes and immune pathways.

## Conclusion

The present study showed that in response to the recent emergence of an epizooty inducing a strong selective pressure, oyster populations generate heritable phenotypic variants that have selective advantage both at the genetic and epigenetic levels. While our study confirms the essential role played by genetics, and also shows that epigenetic variation is strongly controlled by genetic information, we further demonstrate that epigenetic variation can also function independently and, in our case, play a major role in explaining phenotypic variance, with all these mechanisms acting together to rapidly encode new adaptive phenotypes.

## Material and Methods

### Oyster sampling strategy: farming and non-farming areas

Juvenile oysters of *C. gigas* (∼14 old month) were collected in 2018 from six natural populations referred to as B1 to B6 located in the Rade de Brest (France). A total of 356 oysters were collected with ∼60 individuals per population (**Fig. 1A** and **Table S1**). Two populations (B5 and B6) were in oyster “Farming areas” (typical oyster bed with hundreds individuals/m^2^; annual POMS events ^23^), and the four other populations (B1 to B4) were in “Non-farming areas” (> 20 individuals/m^2^; no POMS events). This sampling design enabled the collection of individuals from contrasting environments regarding POMS exposure with “Non-farming” and “Farming areas” hosting high proportion of susceptible and resistant oysters, respectively (**Fig. 1A**). All individuals were brought to Ifremer facilities in Palavas-les-Flots (Montpellier, France) and acclimatized in 45 L tanks for 14 days. In each tank, the seawater temperature was increased by 1°C/day from sampling site temperature to 21°C, continuously UVC-filtered (BIO-UV) and renewed (30%/h). During acclimatization period, all populations were kept in isolated tanks and fed *ad libitum* with Shellfish Diet® 1800 Feeds (Reed Mariculture Inc.).

### Experimental infection by cohabitation between donors and recipient oysters

To classify each oyster as resistant or susceptible phenotype, we performed an experimental infection mimicking POMS event. For this purpose, we used a randomized complete block design composed of eight tanks of 45 L (replicates). Tanks were placed in a water bath to maintain the temperature at 21°C (chiller/heater apparatus; AQUAVIE ICE 3000). In each tank, a water pump (Aquarium System, Maxijet 1000 L/h) and air bubbling maintained water motion and O_2_ level at saturation. Salinity was adjusted to 35 g/L.

To mimic POMS, a cohabitation protocol was used as described ^42^. This started with the injection of 200 µL of OsHV-1 µVar suspension (6.0E7 genomic units) into the adductor muscle of pathogen-free donor oysters. Donor oysters will develop the disease and transmit it through the natural infectious route to the recipient oysters (B1 to B6 populations) (**Fig. 1B**). The ratio between donor and recipient oysters was 1:1 in each tank. The donor oyster populations were composed of 50% of the POMS-susceptible oyster referred family H12 ^17^ and 50% of a genetically diversified standardised oyster spats referred to as NSI ^43^.

Immediately after the OsHV-1 µVar injection, donors and recipient oysters were equally distributed in eight experimental tanks. Disease progression in donor oysters (moribund *vs*. alive) was monitored twice a day (dead donor oysters were removed) during the first 192 hpc after which they were removed from all experimental tanks. Disease progression in recipient oysters started at 24 hpc, and was performed every two hours for 360 hours (no mortalities occurred after day 14).

An oyster was classified as moribund when it could not close its valves after 30 seconds of emersion. Oyster collection at a moribund (susceptible) status enabled the sampling of susceptible oysters before death to avoid DNA degradation. Resistant oysters corresponded to the individuals that were still alive at the end of the experiment (when no death was recorded for 48 hours in all eight tanks). This phenotype was further coded either as a binary trait with the values 0 and 1 corresponding to susceptible and resistant individuals, respectively, or as a semi-quantitative trait corresponding to the survival time (expressed in hours) of an individual after its exposure to the OsHV-1 µVar (i.e. the whole duration of the experiment for the resistant oysters) (**Data S1**). Upon collection, flesh of susceptible and resistant oysters was immediately snap-frozen in liquid nitrogen and stored at -80°C until DNA extraction.

### Survival analysis

Differences in oysters’ survival among the six populations were investigated by Kaplan Meyer model with the ‘*survfit’* and ‘*ggsurvplot’* function of the *survival* (v3.2-11) and *survminer* (v0.4.9) packages, respectively ^44,45^. Cox proportional hazard model was performed using the ‘*coxph’* function from *survival* and was plotted by ‘*ggforest’* function from the *survminer* package. Results were considered significant below the 5% error level.

### Viral load quantification (OsHV-1 µVar)

During the first 192 hpc, 1 mL of water from each experimental tank was sampled daily for viral load quantification. The OsHV-1 µVar DNA was extracted from 200 µL of water using the QIAmp DNA mini Kit (QIAGEN). Quantitative PCR was performed with 5 µL of DNA as described ^46^.

### DNA extraction

Oysters flesh was grounded in liquid nitrogen using 50 mL stainless steel bowls and 20-mm-diameter grinding balls with a vibrational frequency of 30 oscillations per second for 30 seconds (Retsch MM 400 mill). The resulting powder was used for DNA extraction using the NucleoSpin® Tissue kit following manufacturer instructions (MACHEREY-NAGEL GmbH & Co. KG). DNA quantity and purity were checked with a Nanodrop One spectrophotometer (Thermo Scientific) and quality was checked by 0.8% agarose gel electrophoresis. The extracted DNA was stored at -20°C.

### Exome capture and sequencing

The *C. gigas* exome was captured using the SeqCap Epi Enrichment System protocol (Roche Sequencing Solutions, Inc.) ^47^. To capture exons, probes complementary to the whole exonic regions were developed from the genome V9 of *C. gigas* ^48^ in collaboration with Roche. For optimal coverage of the 5’ and 3’ ends of each exon, the probes were designed to cover the 100 base pairs (bp) upstream and downstream to each exon start/end coordinates (**Data S15**).

Exome capture of bisulfite-converted libraries was done according to manufacturer instructions ^47^. Briefly, for each oyster, genomic DNA fragmentation was performed on 1 µg of DNA in addition with DNA phage lambda (GenBank Accession NC_001416) as a spike-in control for the bisulfite conversion efficiency quality control. DNA fragments of average 200 bp were obtained by sonication with the Covaris S220 apparatus (Covaris, Inc.) using the following parameters (Peak Incidence Power: 175, Duty factor: 10, Cycle/Burst: 200, Duration: 70 seconds). After end repair and A-tailing, methylated indexed adapters were ligated and 20 µL of cleaned DNA fragments were subjected to sodium bisulfite conversion using the EZ DNA Methylation-Lightning Kit (Zymo Research, CA). Pre-amplification of the bisulfite-converted library was carried out for an equimolar pool of eight samples and was then subjected to exome capture at 47°C for 45 hours. After cleaning, a final post-capture PCR amplification was performed and captured bisulfite-converted libraries were sequenced (30X) with an Illumina NextSeq 550 system (PE 2 × 150 bp) or an Illumina NovaSeq S1 6000 system (PE 2 × 100 bp).

### SNP and DNA methylation calling

Raw reads quality was checked with FastQC (v0.53) ^49^. Adapter trimming and quality filtering were done with TrimGalore (v0.4.0) ^50^. Bisulfite conversion efficiency was estimated with BSMAPz (v1.1.3) ^51^. BSMAP (v2.90) ^52^ was used to align reads to reference genome V9 of *C. gigas* ^48^. Duplication due to PCR or overlapping between forward and reverse reads were removed following six successive steps (**Fig. S5**): (1) Reads were split into four sets as the top (++ and +-) and the bottom strands (-+ and --) using the ‘split’ option from the BamTools (v1.0.14) ^53^. (2) The top (++ and +-) and bottom (-+ and --) strands were merged to produce a set of top and bottom reads using the ‘merge’ function from BamTools. (3) The top and bottom reads were sorted using the ‘sort’ function from SAMtools (v1.9) ^54^; (4) The PCR duplicates were removed with ‘MarkDuplicates’ function from Picard (v2.21.1) ^55^; (5) The top and bottom read sets were merged back using the ‘merge’ option from BamTools; and (6) The overlapping read pairs were clipped using ‘clipOverlap’ function from bamUtil (v1.0.14) ^56^. The scripts are provided (**Supplementary Information**).

To maximize the accuracy of SNP calling, we used a combination of two SNP callers, FreeBayes (v1.3.1) dedicated to SNP calling from population data ^57^ and MethylExtract (v1.9) dedicated to SNP calling from bisulfite converted sequences ^58^. FreeBayes was first used to call the SNPs present in the dataset including those due to the bisulfite conversion (parameters: --use-best-n-alleles = 2, --use-mapping-quality, --no-partial-observations, -- min-repeat-entropy = 1). Second, MethylExtract was used to call SNPs that were not due to the bisulfite conversion (C/T SNP; parameters: minQ = 20, minDepthSNV = 8, methNonCpGs = 0.9, maxStrandBias = 0.7, varFraction = 0.1, maxPval = 0.05). Finally, only the SNPs identified by both callers were kept and used for Genome-Wide Association Study (GWA) mapping (**Supplementary Information**).

DNA methylation calling in the CG context (CpGs) was done using MethylExtract (same parameters as mentioned above). The methylation data of all samples were combined and used for Epigenome-Wide Association (EWA) mapping (**Supplementary Information**).

### Genome-and Epigenome-wide Association Quality Control (QC)

According to the best practices for GWA mapping ^59^, the following filtering criteria were applied under the PLINK environment (v1.9) ^60^: (1) SNPs supported by a coverage of 8X to 150X were kept; (2) SNPs and samples with a level of missing data above 5% were discarded; (3) SNPs with a minor allele relative frequency (MARF) below 0.05 were discarded; (4) SNPs displaying a significant deviation from Hardy–Weinberg equilibrium (HWE) in resistant (HWE P < 1 × 10E^−06^) and susceptible oysters (HWE P < 1 × 10E^−10^) were excluded; (5) samples with standard deviation of 3 units from mean heterozygosity rate of all samples were discarded; and (6) if present, closely related individuals were excluded to remove cryptic relatedness.

For EWA mapping, the following criteria were performed: (1) CpGs supported by a coverage of 8X to 150X were kept; (2) CpGs and samples with missing data level above 5% were discarded.

For both datasets, the absence of genetic and epigenetic structure between the six populations was checked using an analysis of multivariate homogeneity of group dispersions with the ‘*betadisper*’ function of the *vegan* package (v2.5-7) ^61^. Hierarchical clustering analysis (Euclidian method) and permutational multivariate analysis of variance (PERMANOVA) were performed using the ‘*adonis*’ function from the *vegan* package.

### Genome-and Epigenome wide association mapping

GWA mapping was performed by associating SNPs to the binary trait (using a chi-square allelic test with 1 degree of freedom) and the semi-quantitative trait (using an asymptotic version of Student’s t-test) under the PLINK environment (v1.9) ^60^. EWA mapping was performed by associating DNA methylation variation at each CpGs with the binary and semi-quantitative traits using a linear regression t.test (‘*cpg*.*assoc*’ function) from *CpGassoc* package (v2.60) ^62^. For both GWA and EWA mappings, the significant level of association was defined with a False Discovery Rate (FDR) of 0.05. The GWA/EWA mapping results were visualized using Quantile-Quantile plots and Manhattan plots were generated with *qqman* package (v0.1.8) ^63^. To benefit from a new reference genome assembly with a chromosomal anchor ^64^, homemade scripts were used to positions and visualize SNPs and CpGs on chromosomes (**Supplementary information**).

### Gene Ontology enrichment analysis (GO-terms)

SNPs and CpGs significantly associated with phenotypic variation were subjected to Gene Ontology term enrichment to identify Biological Processes (BP) enriched (**Data S16**). This was done using a Rank-based Gene Ontology Analysis with Adaptive Clustering following the GO_MWU protocol ^65^. The continuous measure of significance used was–log(*P*-value). The following parameters were used for the adaptive clustering: largest = 0.4, smallest = 10, clusterCutHeight = 0.25. A GO-term was considered enriched when FDR < 0.05. We used the REVIGO tool ^66^ to visualise significant GO-terms in a semantic similarity relationship obtained from GO_MWU outcome, with the Uniprot database and a aggregation value of 0.7 with SimRel as semantic similarity measure.

### Genetic and epigenetic correlation and association

Correlative (Mantel test) and association MethQTL approaches between genetic and epigenetic variation were adopted to investigate their relationships. Based on the Spearman correlation coefficient, Mantel test was applied to estimate the correlation between the genetic and epigenetic matrices of dissimilarity among samples. The association between SNPs and CpG methylation levels was identified using a linear regression implemented in *GEM* package (v1.24.0) ^67^ according to the following ‘*Gmodel’*: lm (G ∼ M + covariate), where G and M are the genetic and the DNA methylation level matrix and covariate is the phenotypic trait. This model was applied with the binary and the semi-quantitative trait separately.

### Genetic and epigenetic variation partition

Variation partitioning is a method of using the coefficient of determination to fraction the variation of a response variable into four explanatory variables ^68^. Two of them correspond to the fractions of variance exclusively explained by one of the two explanatory matrices (e.g. genetic or epigenetic), one corresponds to the fraction of variance shared by the two explanatory matrices (i.e. genetic and epigenetic) and the last one corresponds to the fraction of the variance unexplained by the model (**Supplementary Information**).

To estimate the relative contribution of genetic and epigenetic variation to phenotypic variation we used a method developed by ^33^ and applied in ^69^. Briefly, the genetic and epigenetic variance was surrogated by producing Principal Components Analyses (PCA) on the same datasets that were used for GWA/EWA mapping using the ‘*prcomp*’ function (v2.0.0.) *stats* package. Then, using a forward selection method implemented in the ‘*ordistep’* function in the *vegan* package (v2.5-7) ^61^, the best models explaining variance for the binary and semi-quantitative traits were separately obtained with genetic and epigenetic Principal Components (PCs), resulting in four independent models: 2 phenotypic traits × 2 genomic/epigenomic PCs. For each phenotypic trait, the selected PCs from genetic and epigenetic models were retrieved and analysed in a variation partitioning analysis using the ‘*varpart’* function from the *vegan* package (v2.5-7) ^61^.

## Data availability

Raw reads are available at ENA under the project accession number PRJEB60400.

## Author contribution

**GM, DDG, JVD** granted the study

**JDL, YG, BP, GM, CG, JVD** designed the study.

**JDL, YG, FL, JME, BP, BM, LD, JVD** sampled or produced the biological material

**JG, JDL, BM, LD, YG, MS, DDG, CM, ML, PH, CC, CG, JVD** performed the experiments

**JG, JDL, CM, AV, ML, FR, CG, CC, JVD** analyzed the data

**JG, AV, GM, FR, DDG, JVD** wrote the manuscript.

All the authors have revised and approved the manuscript submission

## Acknowledgments

Authors are grateful to the staff of the Ifremer platform of Argenton, Bouin and Palavas-sur-Mer for technical support in animal sampling and experimentation. We are grateful to Dr. Clémentine Vitte and Dr. Benoit Pujol for fruitful discussions, Dr. Yann Dorant for illustrating support and Dr. Guy Riddihough (Life science editors) for his help in manuscript edition. Data used in this work were partly produced through the GenSeq technical facilities. We thank the Bio-Environment platform (University of Perpignan Via Domitia) and Jean-François Allienne for support in sequencing. We thank the bioinformatic service of Ifremer (SEBIMER) for their help in bioinformatics. The present study was supported by the ANR projects DECIPHER (ANR-14-CE19-0023) and DECICOMP (ANR-19-CE20-0004), by the FEAMP project GESTINNOV (grant n°PFEA470020FA1000007), by the EU funded project VIVALDI (H2020 program, n°678589) and by Ifremer (grant politique de site GEM). This study is set within the framework of the “Laboratoires d’Excellences (LABEX)” TULIP (ANR-10-LABX-41).

