## Supplementary for "Epigenetic then genetic variations underpin rapid adaptation of oyster populations (*Crassostrea gigas*) to Pacific Oyster Mortality Syndrome (POMS)": Gawra_et_al_BioRxiv_Supplementary_Information.docx

**Rapid adaptation of oyster populations (*Crassostrea gigas*) to Pacific Oyster Mortality Syndrome (POMS) infectious diseases relies on genetic and epigenetic variations**

Janan Gawra^1^, Alejandro Valdivieso^2^, Julien de Lorgeril^2,3^, Yannick Gueguen^2,4^, Martin Laporte^5^, Fabrice Roux^6^, Mathilde Saccas^2^, Jean-Michel Escoubas^2^, Caroline Montagnani^2^, Delphine Destoumieux-Garzόn^2^, Franck Lagarde^4^, Marc A. Leroy^2^, Philippe Haffner^2^, Bruno Petton^8^, Céline Cosseau^1^, Benjamin Morga^7^, Lionel Dégremont^7^, Guillaume Mitta^1,9^, Christoph Grunau^1^ , Jeremie Vidal-Dupiol^2^*

1 IHPE, Univ Perpignan Via Domitia, CNRS, Ifremer, Univ Montpellier, Perpignan, France

2 IHPE, Univ Montpellier, CNRS, Ifremer, Univ Perpignan Via Domitia, Montpellier, France

3 Ifremer, IRD, Université de la Nouvelle-Calédonie, Université de La Réunion, ENTROPIE, Nouméa, Nouvelle- Calédonie, France

4 MARBEC, Univ Montpellier, CNRS, Ifremer, IRD, Sète, France

5 Division de l'expertise sur la faune Aquatique, Ministère des Forêts, de la Faune et des Parcs (MFFP), 880 chemin Sainte-Foy, G1S 4X4, Québec, Québec, Canada

6 LIPME, INRAE, CNRS, Université de Toulouse, Castanet-Tolosan, France

7 Ifremer, ASIM, Adaptation Santé des Invertébrés Marins, La Tremblade, France

8 LEMAR UMR 6539, UBO/CNRS/IRD/Ifremer, 11 presqu’île du vivier, 29840 Argenton-en-Landunvez, France

9 Univ Polynesie Francaise, ILM, IRD, Ifremer, F-98719 Tahiti, French Polynesia, France

Keywords: rapid adaptation, genetic, epigenetic, POMS, oyster, exome capture

**INDEX**

**Supplementary tables, figures, and information**

**Tables**

Supplementary table 1 …..……………………………………………………………………….3

Supplementary table 2 …..……………………………………………………………………….4

Supplementary table 3 …..……………………………………………………………………….5

**Figures**

Supplementary figure 1…..……………………………………………………………………….6

Supplementary figure 2…..……………………………………………………………………….7

Supplementary figure 3…..……………………………………………………………………….8

Supplementary figure 4…..……………………………………………………………………….9

Supplementary figure 5…..……………………………………………………………………..10

**Information**

Supplementary information ……………………………………………………………..11-28

### **Supplementary Tables**

#### **Table S1**. The coordinates and the number of oysters selected in each population

| **Area** | **Population** | **Latitude** | **Longitude** | **Number of oysters** |
| --- | --- | --- | --- | --- |
| Non-farming | B1 | 48.379364 | -4.446286 | 61 |
|  | B2 | 48.341789 | -4.441086 | 59 |
|  | B3 | 48.322392 | -4.454078 | 61 |
|  | B4 | 48.296575 | -4.451778 | 56 |
| Farming | B5 | 48.32815 | -4.321947 | 59 |
|  | B6 | 48.34695 | -4.338986 | 60 |
|  |  |  | Total | 356 |

#### **Table S2.** Hazard ratios of the relative risk of mortality for all six oysters populations.

| **Area** | **Population** | **Number of oysters** | **Hazard ratio (95%, CI)** | ***P*-value** |
| --- | --- | --- | --- | --- |
| Non-Farming | B1 | 61 | Reference * | NA |
|  | B2 | 59 | 0.595 (0.3732-0.95) | *0.029* |
|  | B3 | 61 | 10550 (0.6840-1.61) | 0.825 |
|  | B4 | 56 | 0.767 (0.4831-1.22) | 0.261 |
| Farming | B5 | 59 | 0.046 (0.0143-0.15) | *< 0.001* |
|  | B6 | 60 | 0.030 (0.0072-0.12) | *< 0.001* |

* B1 is the reference

#Events: 150; Global *P*-value (Log-Rank): 2.2141e^-27^

AIC: 1563.55; Concordance index: 0.74

#### **Table S3.** Methylation Quantitative Trait Loci (MethQTL) using the covariate as binary or semi-quantitative trait associate with Pacific Oyster Mortality Syndrome (POMS)

| **MethQTL** | **Binary** | **Semi-quantitative** |
| --- | --- | --- |
| **Total SNPs** | 214,263 | 214,263 |
| **Total CpGs** | 635,201 | 635,201 |
| **Significant SNP-CpG pairs** | 5,151,194 | 5,152,611 |
| **Total non-redundant SNPs** | 160,325 | 160,22 |
| **Total non-redundant CpGs** | 557,703 | 557,85 |
| **MethQTL controlling a CpG identified in EWA mapping** | 207 | 198 |
| **MethQTL identified by the GWA mapping (suggestive threshold)** | 3 | 8 |
| **CpGs identified by the EWA mapping and controlled by a MethQTL** | 126 | 111 |
| **MethQTL involving a CpG and a SNP identified by the EWA and GWA mapping** | 18 | 15 |

### **Supplementary Figures**

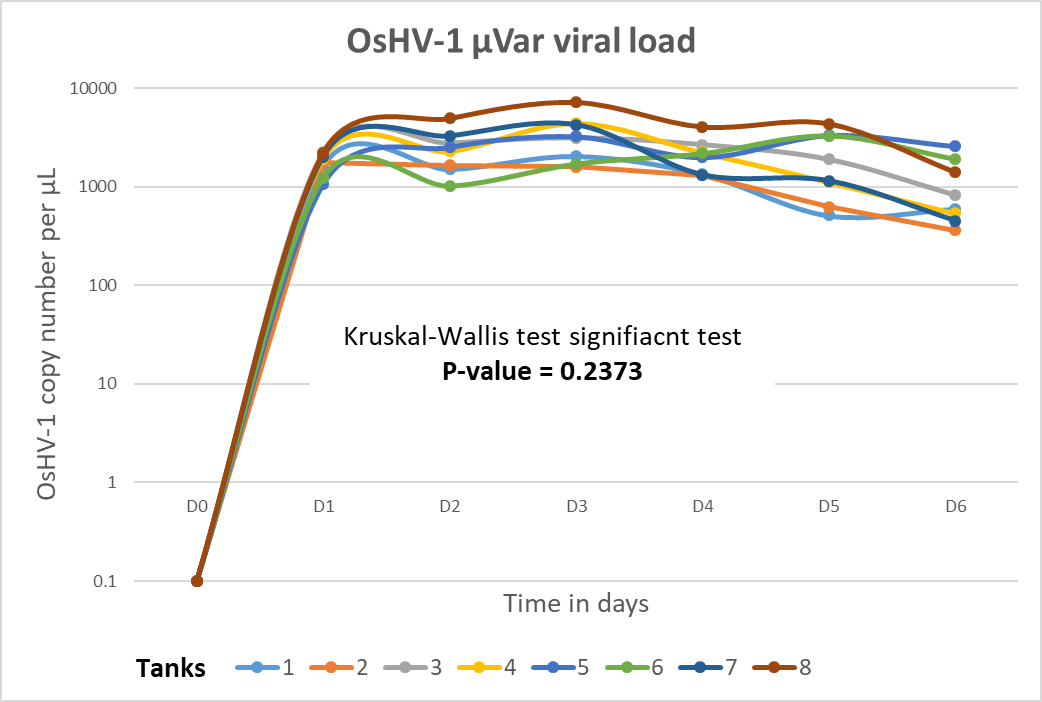

#### **Figure S1.** The OsHV-1 μVar viral load from the seawater was taken in the eight tanks during the first seven days of cohabitation.

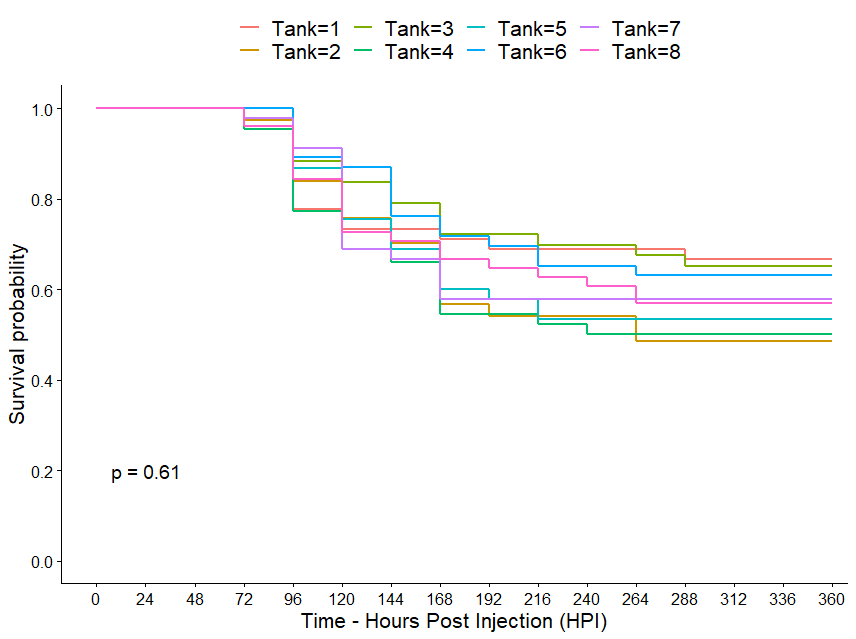

Survival probability

Hours post-cohabitation (hpc)

#### **Figure S2.** The Kaplan–Meier survival curves of oysters among the eight experimental tanks.

Dimension 2 (0.55%)

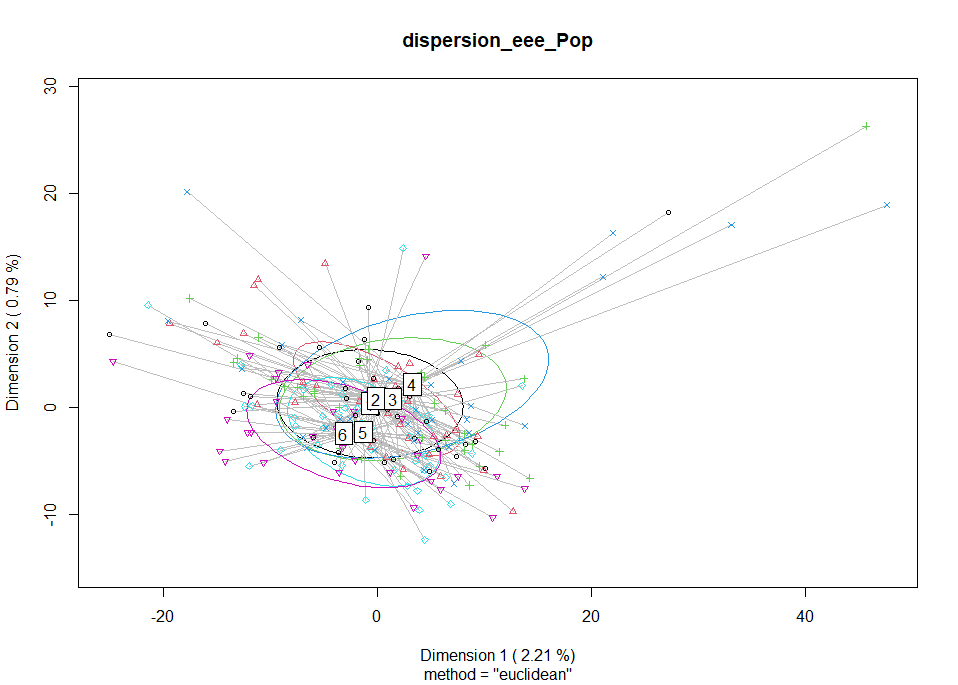

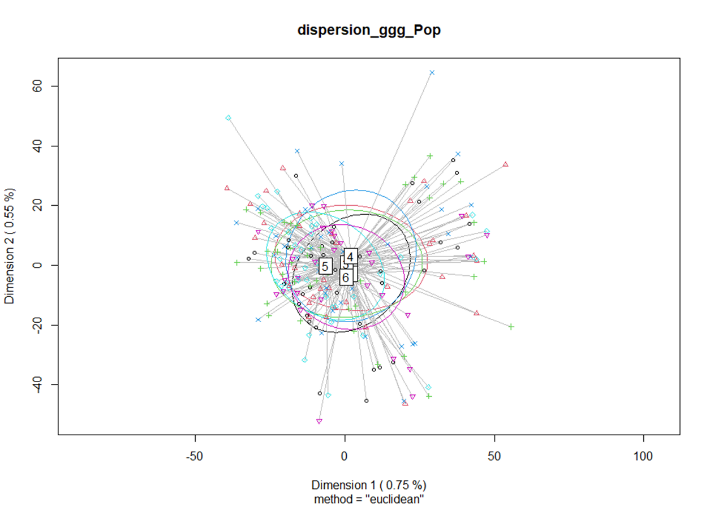

Dimension 2 (0.79%)

Dimension 1 (0.75%)

Dimension 1 (2.21%)

A

B

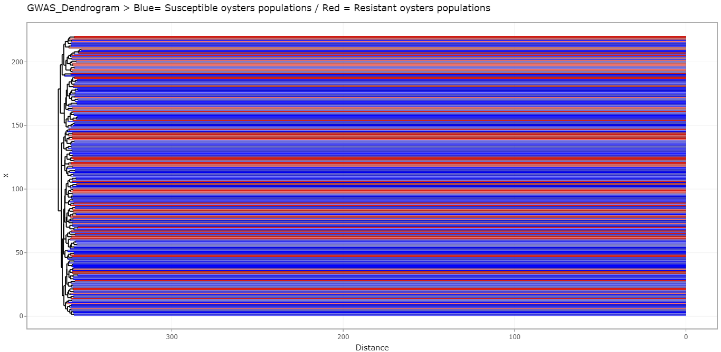

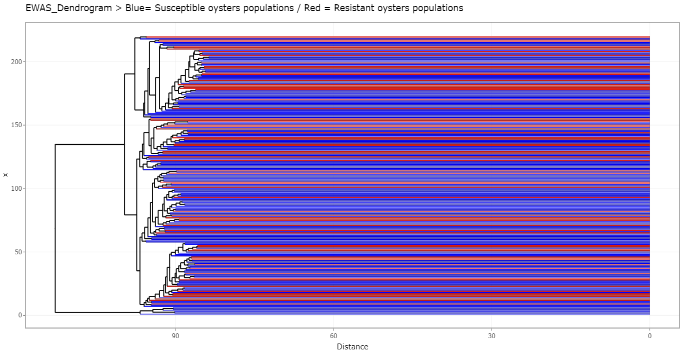

Genetic

Epigenetic

C

D

`

Distance

Distance

#### **Figure S3.** Analysis of multivariate homogeneity of group dispersions (variances) among the six populations for **A)** Genetic and **B)** Epigenetic data. Hierarchical cluster analysis for the **C)** Genetic and **D)** Epigenetic data. The blue and red gradient color corresponds to the “Non-farming” and “Farming” populations.

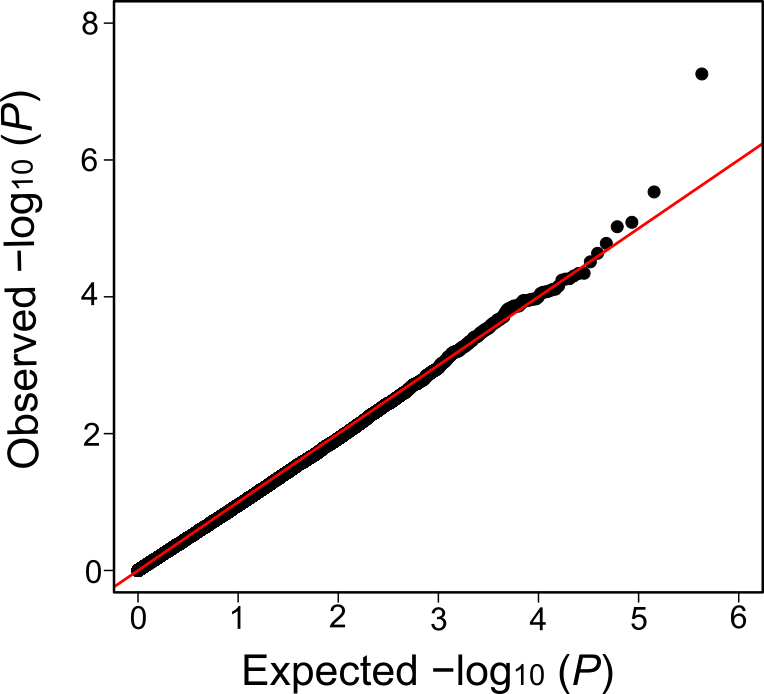

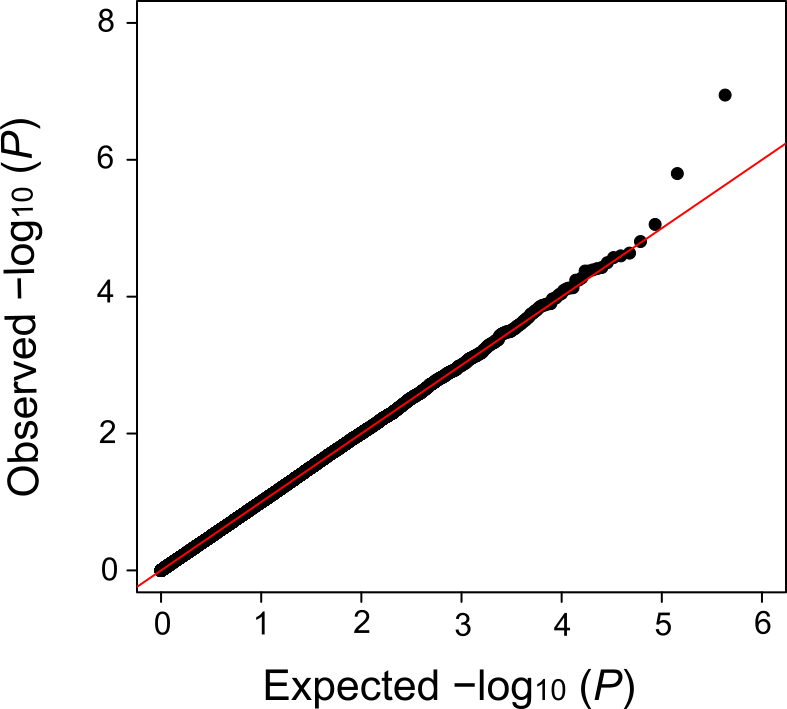

**A**

**B**

Binary

Semi-quantitative

SNPs

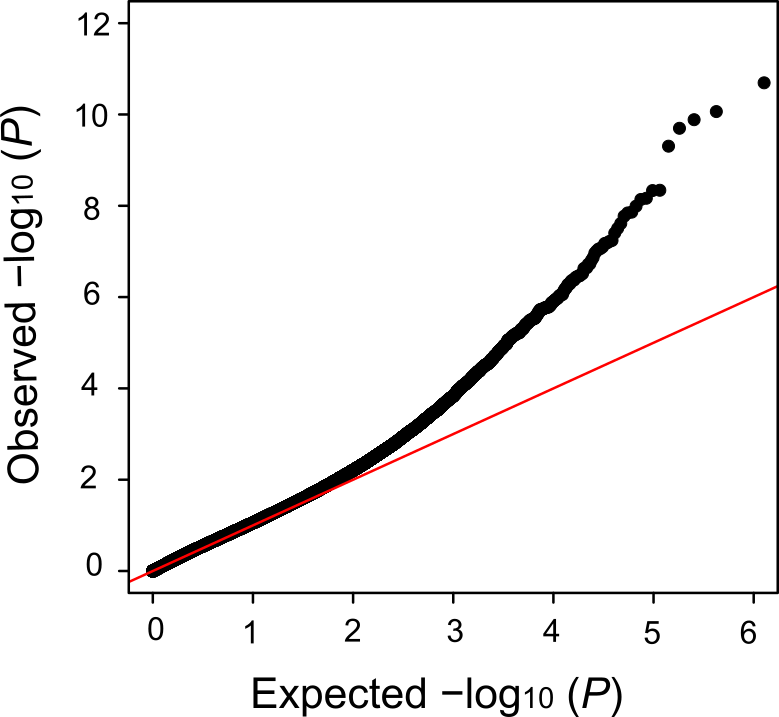

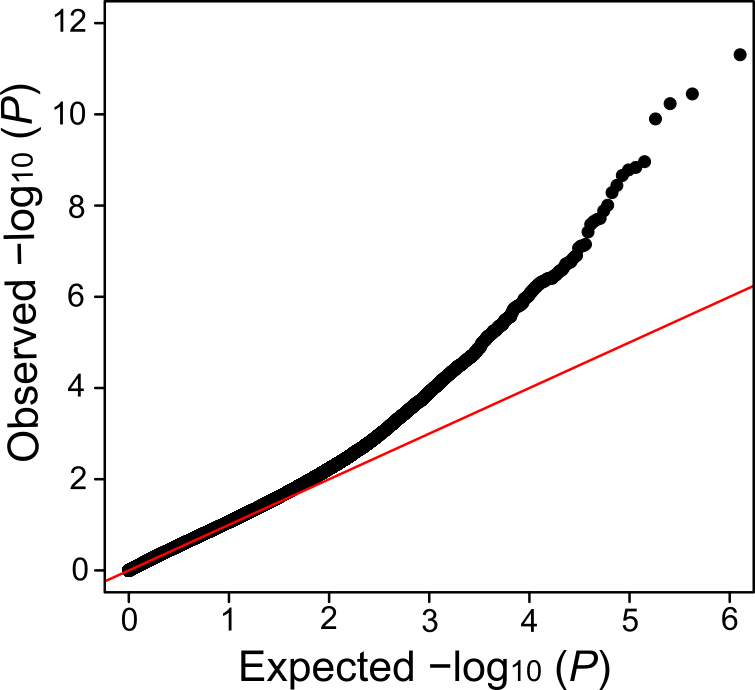

CpGs

**C**

**D**

#### **Figure S4.** The quantile-quantile (Q-Q plot) of the significant SNPs identified in A) binary and B) semi-quantitative trait, and Q-Q plot of the significant CpGs in C) binary and D) semi-quantitative trait.

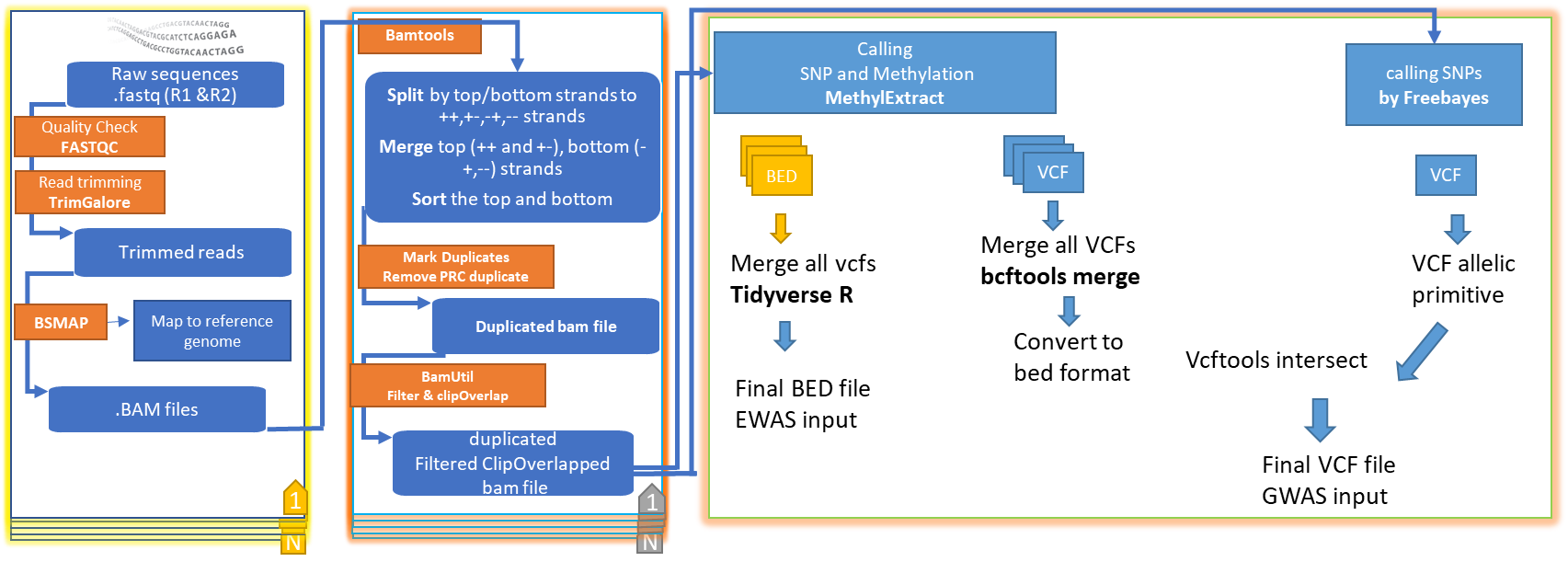

#### **Figure S5.** Bioinformatics pipeline steps applied for the identification of SNP and DNA methylation callings. See below for bioinformatics scripts.

### **Supplementary Information**

Bioinformatic scripts

#### SNP and DNA methylation Calling

Before the first step - trimming the data to remove the adapter and quality check.

The parameters for trimming: For 150 bp, 2x Paired End reads [remove the last 50]

trim_galore --paired --illumina --quality ${params.quality} --three_prime_clip_R1 50 --three_prime_clip_R2 50

\##### While for trimming reads with 100 bp 2x Paired End reads ####

trim_galore --paired --illumina --quality ${params.quality} --clip_R1 1 --clip_R2 1

Then to do the Mapping - filtering and SNP - Methylation calling

The pipeline for this is ready. On gitlab Ifremer

<https://gitlab.ifremer.fr/bioinfo/nf-core-gem.git>

Note is better to select only samples that have a closely number of reads otherwise it will lead to many missing data in the VCF (SNP) and Bed (DNA methylation) files

Step 1 - Map reads to reference genome using bsmap

bsmap -r ${params.bsmap_repeat} -n ${params.bsmap_mapstrand} -s ${params.bsmap_seedsize} -p ${task.cpus} -d ${params.genome} -a ${name}_R1_val_1.fq -b ${name}_R2_val_2.fq -o ${name}.sam &> bsmap-${name}.log 2>&1

picard -Xms512m -Xmx${task.memory.toGiga()}g AddOrReplaceReadGroups RGID=${name} RGLB=${name} RGPL=illumina RGSM=${name} RGPU=@A00902:117:HKKNJDRXX:2 VALIDATION_STRINGENCY=LENIENT I=${name}.sam O=${name}.bam &> picard-${name}.log 2>&1

STEP 2 - Split, merge and sort mapped reads using bamtools

bamtools split -tag ${params.bamtools_tag} -in ${bam} &> bamtools-${name}.log 2>&1

bamtools merge -in ${name}.TAG_${params.bamtools_tag}_++.bam -in ${name}.TAG_${params.bamtools_tag}_+-.bam -out ${name}_top_merged.bam &>> bamtools-${name}.log 2>&1

bamtools merge -in ${name}.TAG_${params.bamtools_tag}_-+.bam -in ${name}.TAG_${params.bamtools_tag}_--.bam -out ${name}_bottom_merged.bam &>> bamtools-${name}.log 2>&1

samtools sort ${name}_top_merged.bam > ${name}_top_merged_sorted.bam 2> samtools-${name}.log

samtools sort ${name}_bottom_merged.bam > ${name}_bottom_merged_sorted.bam 2>> samtools-${name}.log

STEP 3 - Mark duplicates with picard tools - remove duplicates

picard -Xms512m -Xmx${task.memory.toGiga()}g -Djava.io.tmpdir=./picard MarkDuplicates \

VALIDATION_STRINGENCY=SILENT \

INPUT=${topbam} \

OUTPUT=${name}_top_rm_dupl.bam \

METRICS_FILE=${name}_top_rm_dupl_metrics.txt \

ASSUME_SORTED=TRUE \

REMOVE_DUPLICATES=TRUE \

CREATE_INDEX=TRUE &> ${name}_top_picard.log 2>&1

picard -Xms512m -Xmx${task.memory.toGiga()}g -Djava.io.tmpdir=./picard MarkDuplicates \

VALIDATION_STRINGENCY=SILENT \

INPUT=${bottombam} \

OUTPUT=${name}_bottom_rm_dupl.bam \

METRICS_FILE=${name}_bottom_rm_dupl_metrics.txt \

ASSUME_SORTED=TRUE \

REMOVE_DUPLICATES=TRUE \

CREATE_INDEX=TRUE &> ${name}_bottom_picard.log 2>&1

STEP 4 - Merge reads with bamtools

bamtools merge -in ${topbam_rm_dupl} -in ${bottombam_rm_dupl} -out ${name}_bsmap_non_masked_rm-dupl.bam &> bamtools-${name}.log 2>&1

STEP 5 - Filter merged reads with bamtools

bamtools filter \

-isMapped true \

-isPaired true \

-isProperPair true \

-forceCompression \

-in ${merged_bam} \

-out ${name}_filtered.bam &> bamtools-${name}.log 2>&1

STEP 6 - Filter clipoverlap with bamutils and index bam file

bam clipOverlap \

--stats \

--in ${filtered_bam} \

--out ${name}_clipped.bam &> bamtutils-${name}.log 2>&1

STEP 7 - Methylation maps and SNP calling with MethylExtract

MethylExtract.pl p=${task.cpus} seq=${params.genome} inDir=. outDir=. minDepthMeth=${params.methylextract_mindepthmeth} minDepthSNV=${params.methylextract_mindepthsnv} context=ALL wigOut=Y bedOut=Y flagW=${params.methylextract_flagw} flagC=${params.methylextract_flagc} &> ${name}_methylextract.log

Then is merge the Bed files AND VCF files

Two problems here to deal with. That maybe come from the not well having a homogenous number of reads, if a closely number of reads have selected maybe these problem probably won’t show up.

First: merge the vcf files (Single Nucleotide Polymorphisms; SNPs containing file)

Because with the METHYLEXTRACT package the SNP calling is done based on single sample producing single vcf file reporting only the SNPs. When merging many VCF files from different sample would lead to many missing data. Simply because one sample or many samples would have a homozygote genotype for reference allele and others would have a heterozygote or homozygote genotype for alternative allele.

**How to tackle this issue?**

First, we use FreeBayes to obtain a single VCF file for all the samples.

Script for FREEBAYES

First, we use FreeBayes to obtain a single VCF file.

#PBS -q omp

#PBS -l walltime=120:00:00

#PBS -l mem=115g

#PBS -l ncpus=56

\## Manage script history

INPUT_DIR=/home/datawork-ihpe/gem/06_clipped-bam-files

FreeBayes_TOOLS=". /appli/bioinfo/freebayes/latest/env.sh"

#GENOME=/home1/datawork/jgawra/GWAS_EWAS_TEST/vcfcd $INPUT_DIR

$FreeBayes_TOOLS

########################################

\## Shell variables ##

########################################

#INPUT_DIR=/home1/datawork/jgawra/GWAS_EWAS_TEST

GENOME=/home1/datawork/jgawra/GWAS_EWAS_TEST/vcf/oyster.v9.fa

OUTPUT_DIR=/home1/scratch/jgawra/GWAS_EWAS_TEST

########################################

\## prepapre input file ##

########################################

ls -d "${INPUT_DIR}/"*"_clipped.bam" > "${INPUT_DIR}/SAMPLES_clipped_bam.txt"

########################################

\## Freebays variant calliing ##

########################################

echo "create a list of bam files... "`cat "${INPUT_DIR}/SAMPLES_clipped_bam.txt"`

echo "variant calling..."

freebayes-parallel <(fasta_generate_regions.py "${GENOME}.fai" 10000) 56 -p 2 -f $GENOME --use-best-n-alleles 2 --use-mapping-quality --min-coverage 8 --no-partial-observations --min-repeat-entropy 1 -L ${INPUT_DIR}/SAMPLES_clipped_bam.txt >& ${OUTPUT_DIR}/freebayes_248_Brest_samples_s

econd_try.vcf 2> ${OUTPUT_DIR}/freebayes_248_Brest_samples_second_try.vcf.log

Then I need to check if the vcf file is ok. Then is to change the vcf from the haplotype to single SNP type by using the below script

#!/usr/bin/env bash

#PBS -q omp

#PBS -l walltime=250:00:00

#PBS -l mem=115g

#PBS -l ncpus=56

DATA=/home/datawork-ihpe-gem-nos

BCFTOOLS_TOOLS=". /appli/bioinfo/vcflib/1.0.0_rc1/env.sh"

cd $DATA

$BCFTOOLS_TOOLS

vcfallelicprimitives -kg freebayes_248_Brest_samples_second_try_remove-bad-lines.vcf > freebayes_248_Brest_samples_second_try_remove_bad_lines_vcfallelicprimitives.vcf

FreeBayes is not able to differentiate a real SNP fro independently in separate VCF files for each sample re, MethylExtract was used to call real SNPs in separate VCF files for each sample independently. This is done in STEP 7 already [page 4].

Then, we used BCFTOOLS to merge all the VCF files into a single VCF file and convert it to a bed file format (which contains the real SNP genomic location).

#!/usr/bin/env bash

#PBS -q sequentiel

#PBS -l walltime=00:30:00

#PBS -l mem=5g

DATA=/home1/datawork/jgawra/GWAS_EWAS_TEST/vcf/vcf-sub

BCFTOOLS_TOOLS=". /appli/bioinfo/bcftools/latest/env.sh"

cd $DATA

$BCFTOOLS_TOOLS

\#### First is to bgzip the vcf file to index it.

for file in *_sub.vcf

do

bgzip -c $file

done

\#### Then to index it.

#for file in *.vcf.gz ; do bcftools index -c $file ; done

\#### Finally to merge it.

#bcftools merge --force-samples *vcf.gz -Oz -o Merged.vcf.gz

Then is to make a bed file like to of all the vcf position (the real SNPs) that to be used in the next step

Finally, we intersected it with a VCF file produced by FreeBayes using vcfintersect from VCFTOOLS (version 0.1.16) and obtained the final VCF file that was will be used for GWA mapping analyses.

#!/usr/bin/env bash

#PBS -q omp

#PBS -l walltime=50:00:00

#PBS -l mem=50g

#PBS -l ncpus=28

DATA=/home/datawork-ihpe-gem-nos

BED_DATA=/home/datawork-ihpe/gem/06_clipped-bam-files

#BCFTOOLS_TOOLS=". /appli/bioinfo/vcflib/1.0.0_rc1/env.sh"

VCFTOOLS=". /appli/bioinfo/vcftools/latest/env.sh"

cd $DATA

$VCFTOOLS

Vcftools

--vcf freebayes_248_Brest_samples_second_try_remove_bad_lines_vcfallelicprimitives.vcf

--bed ${BED_DATA}/vcf2bed_jb.bed

--out freebayes_248_samples_vcfallelicprimitive_with_region_methylextract.vcf

--temp $SCRATCH –recode

\#### This file is ready for GWA mapping analysis

####### This is a vcf file that would be used for GWA mapping analysis.

Second: How to deal with Bed file

First I used R to merge all the bed files (produced by methyextract in step 7; page 3).

#first I need to prepare the bed files

#!/usr/bin/env bash

#PBS -q omp

#PBS -l walltime=48:00:00

#PBS -l select=1:ncpus=28:mem=115g

DATA=/home/datawork-ihpe/gem/09_methylextract_results/

cd $DATA

for i in *.bed

do

sed '1d' "$i" > ${i%.bed}_temp1.bed

done

for i in *_temp1.bed

do

awk '{print $1,($3 - 1),($5/10)}' "$i" > ${i}_temp2.bed

done

for i in *_temp2.bed

do

sed -i '1i chrom pos methratio' "$i"

done

for i in *_temp2.bed

do

sed -e 's/ */\t/g' "$i" > ${i%CG_temp1.bed_temp2.bed}CG_2.bed

done

Then is to merge all the bed files

Merge <- tibble(chrom = "C13972",

pos = 83,

methratio_ValueUseless = 0)

for (file in list.files(path=".", pattern="*CG_2.bed")	) {

filename <- file

Merge <- full_join(Merge,read_tsv(file) %>%

rename(!!filename := methratio) %>%

as_tibble() )

}

Merged3.bed <- Merge %>% select(-methratio_ValueUseless)

\#### This file is ready for EWA mapping analysis

Locating the SNPs and CpGs in the new released Genome of Crassostrea gigas (Peñaloza et al., 2021).

SNPs locating in the new genome

Locating the SNPs passing the PLINK quality control to the NEW Roslin GENOME (with chromosome information). This is for visualization purpose, so we can have a Manhattan plot with the ten chromosome.

Preparing a FASTA file

\### The final plink output file (binary fileset; that was used for GWA mapping; containing 214,263 SNPs) were mapped to the new genome CGA using the vcfprimer from

\## First the final plink binary fileset were converted to vcf file.

\### Then for each SNP, a 100 bps were added and finally producing a fasta file that each line is a SNP with 100 bps following.

\### Script for making a FASTA file by vcfprimer v1.0.0. on Linux.

#!/usr/bin/env bash

#PBS -q omp

#PBS -l walltime=40:00:00

#PBS -l mem=20g

#PBS -l ncpus=8

##call the vcflib tool

. /appli/bioinfo/vcflib/1.0.0_rc1/env.sh

\## data location

DATA=/home/datawork-ihpe-gem-nos/CGA-genome

cd $DATA

vcfprimers /home/datawork-ihpe-gem-nos/filtered_Depth_vcf_8-150/plink.vcf -f oyster.v9.fa -l 100 > reads-100bp_plink_214k.fasta

Align the fasta file to new genome

\#### Then the fasta file is used to align it on the new genome using the BOWTIE2 v2.3.5

B.1- First we make an index for the genome

#!/usr/bin/env bash

#PBS -q omp

#PBS -l mem=50G

#PBS -l ncpus=28

#PBS -l walltime=10:00:00

BANK_DIR=/home/datawork-ihpe-gem-nos/CGA-genome

BANK_FILE_NAME=/home/datawork-ihpe-gem-nos/CGA-genome/GCA_902806645.1_cgigas_uk_roslin_v1_genomic.fna

\# Le nom de la banque tel qu'on l'utilisera avec Bowtie2

INDEX_NAME=/home/datawork-ihpe-gem-nos/CGA-genome/GCA_902806645.1_cgigas_uk_roslin_v1_genomic

\# Lancement de Bowtie

bowtie2_cpus=$((${NCPUS}-2))

. /appli/bioinfo/bowtie2/2.3.5/env.sh

cd ${BANK_DIR}

bowtie2-build ${BANK_FILE_NAME} ${INDEX_NAME} -p ${bowtie2_cpus} >& $BANK_DIR/mkbowtie.log 2>&1

B.2- Align the fasta to the new genome and produce a SAM file

#!/usr/bin/env bash

#PBS -q omp

#PBS -l mem=50G

#PBS -l ncpus=28

#PBS -l walltime=10:00:00

BANK_DIR=/home/datawork-ihpe-gem-nos/CGA-genome

#BANK_FILE_NAME=/home/datawork-ihpe-gem-nos/CGA-genome/GCA_902806645.1_cgigas_uk_roslin_v1_genomic.fna

\# Le nom de la banque tel qu'on l'utilisera avec Bowtie2

INDEX_NAME=/home/datawork-ihpe-gem-nos/CGA-genome/GCA_902806645.1_cgigas_uk_roslin_v1_genomic

\# launching the BOWTIE2

bowtie2_cpus=$((${NCPUS}-2))

. /appli/bioinfo/bowtie2/2.3.5/env.sh

cd ${BANK_DIR}

bowtie2 -x ${INDEX_NAME} -f ${BANK_DIR}/reads-100bp_plink_414k.fasta -S reads-50bp_plink_214.fasta.sam

Convert the SAM file to Bed file

\#### Then convert the SAM file were converted to bed using Linux cat, grep and sed and awk tools

cat reads-100bp_plink_214.fasta.sam | grep -v "@HD" | grep -v "@SQ" |grep -v "@PG" | awk '{print $3"\t"$1"\t"($4+100)"\t"$5}' | sed 's/_LEFT//g' > LR_reads-100bp_plink_214K.fasta.sam.bed

\### The output would be a SNP, and the position of the SNP in the chromosome and its coordinates.

Intersecting with the GWA mapping output file to be used for Manhattan plot

\#### Then this bed file were intersected with the GWA mapping output using the tidyverse package by left_join function in R.

library(tidyverse)

setwd("E:/GEM/PhD-Thesis/Lab Methodology/GWAS_EWAS_ANALYSIS/plink/Final_GWAS")

\### read the file with the SNPs and their coordinates in the new genome

df <- read.delim("E:/GEM/PhD-Thesis/Lab Methodology/GWAS_EWAS_ANALYSIS/plink/Final_GWAS/LR_reads-100bp_plink_214K.fasta.sam.bed", header=FALSE)

\## load the GWAS association file to intersect it with the bed file to located each SNP in the new genome

assoc_results <- read.csv("E:/GEM/PhD-Thesis/Lab Methodology/GWAS_EWAS_ANALYSIS/plink/Final_GWAS/assoc_results.assoc", sep="")

assoc_results_pq.qassoc <- read.csv("E:/GEM/PhD-Thesis/Lab Methodology/GWAS_EWAS_ANALYSIS/plink/Final_GWAS/assoc_results-pq.qassoc", sep="")

colnames(df)[1] <- "CHR"

colnames(df)[2] <- "SNP"

colnames(df)[3] <- "POS"

colnames(df)[4] <- "Q"

df2<- cbind(df, read.table(text = as.character(df$CHR), sep = '_'))

df2$SNP <- paste(df2$V2,df2$V3, sep="_")

df3 <- df2[c(5,1,2,3)]

library(dplyr)

df4 <- df3 %>%

group_by(SNP) %>%

filter(POS==max(POS))

df5 =df4[!duplicated(df4$SNP), ]

logistic_adjusted_merged <- left_join(df5, assoc_results , by=c("SNP"))

logistic_adjusted_merged <- left_join(df5, assoc_results_pq.qassoc , by=c("SNP"))

colnames(logistic_adjusted_merged)[1] <- "CHR"

logistic_adjusted_merged$CHR [grepl("CADCXH*", logistic_adjusted_merged$CHR)] <- "88"

logistic_adjusted_merged$CHR [grepl("LR761634.1", logistic_adjusted_merged$CHR)] <-"1"

logistic_adjusted_merged$CHR [grepl("LR761635.1", logistic_adjusted_merged$CHR)] <-"2"

logistic_adjusted_merged$CHR [grepl("LR761636.1", logistic_adjusted_merged$CHR)] <-"3"

logistic_adjusted_merged$CHR [grepl("LR761637.1", logistic_adjusted_merged$CHR)] <-"4"

logistic_adjusted_merged$CHR [grepl("LR761638.1", logistic_adjusted_merged$CHR)] <-"5"

logistic_adjusted_merged$CHR [grepl("LR761639.1", logistic_adjusted_merged$CHR)] <-"6"

logistic_adjusted_merged$CHR [grepl("LR761640.1", logistic_adjusted_merged$CHR)] <-"7"

logistic_adjusted_merged$CHR [grepl("LR761641.1", logistic_adjusted_merged$CHR)] <-"8"

logistic_adjusted_merged$CHR [grepl("LR761642.1", logistic_adjusted_merged$CHR)] <-"9"

logistic_adjusted_merged$CHR [grepl("LR761643.1", logistic_adjusted_merged$CHR)] <-"10"

\##

logistic_adjusted_merged$CHR <- gsub("\\*", "99", logistic_adjusted_merged$CHR)

###write.table(logistic_adjusted_merged, file = "ADD_logistic_merged.txt", sep = "\t", quote = FALSE, row.names =F)

logistic_adjusted_merged$CHR<-as.numeric(logistic_adjusted_merged$CHR)

logistic_adjusted_merged$POS<-as.numeric(logistic_adjusted_merged$POS)

logistic_adjusted_merged$P<-as.numeric(logistic_adjusted_merged$P)

colnames(logistic_adjusted_merged)[2] <- "chr_position_dir"

colnames(logistic_adjusted_merged)[5] <- "chr_v9"

write.table(logistic_adjusted_merged, file ="assoc_results_merged.txt", sep = "\t", row.names = FALSE)

write.table(logistic_adjusted_merged, file ="assoc_results_pq.qassoc_merged.txt", sep = "\t", row.names = FALSE)

Plotting the Manhattan plot and QQplot

library(qqman)

\### Manhattan plot

jpeg(filename = "2Manhattan_plot_gwas_binary.jpg", width = 1200, height = 550 )

manhattan(x = assoc_results_merged, chr = "CHR", bp = "POS", p = "P", genomewideline = -log10(0.05/214318), suggestiveline = -log10(0.0005), col = c("blue", "red"))

dev.off()

jpeg(filename = "2Manhattan_plot_gwas_coninous.jpg", width = 1200, height = 550 )

manhattan(x = assoc_results_pq.qassoc_merged, chr = "CHR", bp = "POS", p = "P", genomewideline = -log10(0.05/214318), suggestiveline = -log10(0.0005), col = c("blue", "red"))

dev.off()

####qqplot

jpeg(filename = "qq_plot_gwas_binary.jpg")

qq(assoc_results_merged$P)

dev.off()

jpeg(filename = "qq_plot_gwas_continous.jpg")

qq(assoc_results_pq.qassoc_merged$P)

dev.off()

Intersect the CpG to the new Genome

First prepare bed file

\## First we prepare a bed file. It is a CpG and its start position and end position for a CpG.

CpG_List = results_95_220_max [c(1)]

df2<- cbind(CpG_List, read.table(text = as.character(CpG_List$TargetID), sep = "_"))

CpG_List = df2[c(2,3)]

write.table(CpG_List, file ="CpG_List-635K.txt", sep = "\t",row.names = F, quote = F,col.names = F )

Prepare a fasta file

\### To do this, a getfasta function from bedtools was used to get a fasta file.

awk '{print $1"\t"$2"\t"($2+49)"\t"(49-$2)}' CpG_List-635K.txt > tmp1

sed 's/-//g' tmp1 > tmp2

awk '{print $1"\t"$4"\t"$3}' tmp2 > tmp3

bedtools getfasta -fi oyster.v9.fa -bed tmp3 -name > tmp99

Convert the Fasta to bed file

cat 2getfasta_CpG_filtered_bed.fasta.sam | grep -v "@HD" | grep -v "@SQ" |grep -v "@PG" | awk '{print $3"\t"$1"\t"($4+49)"\t"$5}' | sed 's/_LEFT//g' > 2getfasta_CpG_filtered_bed.fasta.sam.bed

Intersecting the Bed with EWA mapping output

get2<- read.table("E:/GEM/PhD-Thesis/Lab Methodology/GWAS_EWAS_ANALYSIS/cpgassoc/Final_EWAS/2getfasta_CpG_filtered_bed.fasta.sam.bed", quote="\"", comment.char="")

get = get2 [c(1,2,3)]

colnames(get)[1] <- "CHR"

colnames(get)[2] <- "CHR_Pos_Start_End"

colnames(get)[3] <- "MAPINFO"

get$CHR <- as.character(get$CHR)

get$CHR [grepl("CADCXH*", get$CHR)] <- "88"

get$CHR [grepl("LR761634.1", get$CHR)] <-"1"

get$CHR [grepl("LR761635.1", get$CHR)] <-"2"

get$CHR [grepl("LR761636.1", get$CHR)] <-"3"

get$CHR [grepl("LR761637.1", get$CHR)] <-"4"

get$CHR [grepl("LR761638.1", get$CHR)] <-"5"

get$CHR [grepl("LR761639.1", get$CHR)] <-"6"

get$CHR [grepl("LR761640.1", get$CHR)] <-"7"

get$CHR [grepl("LR761641.1", get$CHR)] <-"8"

get$CHR [grepl("LR761642.1", get$CHR)] <-"9"

get$CHR [grepl("LR761643.1", get$CHR)] <-"10"

get$CHR <- as.numeric(get$CHR)

#newthing

get$CHR_Pos_Start_End <- gsub (":", "-", get$CHR_Pos_Start_End)

df2<- cbind(get, read.table(text = as.character(get$CHR_Pos_Start_End), sep = "-"))

df2$TargetID = paste(df2$V1, df2$V2+49,sep="_")

\# colnames(df2)[4] <- "TargetID"

\# colnames(df2)[5] <- "Pos"

\#

\# df2$TargetID <- gsub (":", "_", df2$TargetID)

library(tidyverse)

colnames(results_95_220_max)[1] <- "TargetID"

colnames(results_cpg_death2)[1] <- "TargetID"

merge_binary <- left_join(results_cpg_death2, df2, by="TargetID")

merge_continous<- left_join(results_95_220_max, df2, by="TargetID")

merge_binary$CHR[is.na(merge_binary$CHR)] <- 99

merge_binary$MAPINFO[is.na(merge_binary$MAPINFO)] <- 100

merge_continous$CHR[is.na(merge_continous$CHR)] <- 99

merge_continous$MAPINFO[is.na(merge_continous$MAPINFO)] <- 100

write.table(merge_binary, file ="merge_binary_Ewas.txt", sep = "\t", row.names = FALSE)

write.table(merge_continous, file ="merge_continous_Ewas.txt", sep = "\t", row.names = FALSE)

merge_continous_Ewas <- read.delim("C:/GEM_THESE/Final_EWAS/merge_continous_Ewas_CpG_GENE2.txt")

merge_binary_Ewas <- read.delim("C:/GEM_THESE/Final_EWAS/merge_Binary_Ewas_CpG_GENE2.txt")

colnames(merge_continous_Ewas)[1] <- "SNP"

colnames(merge_binary_Ewas)[1] <- "SNP"

summary(merge_binary_Ewas)

merge_binary_Ewas$P.value[is.na(merge_binary_Ewas$P.value)] <- 1

merge_continous_Ewas$P.value[is.na(merge_continous_Ewas$P.value)] <- 1

library(qqman)

\#### plotting by qqman package

jpeg(filename = "Manhattan_plot_ewas_binary.jpg", width = 1200, height = 550 )

manhattan(x = merge_binary_Ewas, chr = "CHR", bp = "MAPINFO", p = "P.value", genomewideline = -log10(1.901340e-05),suggestiveline = FALSE, col = c("blue", "red"))

dev.off()

jpeg(filename = "Manhattan_plot_ewas_continous.jpg", width = 1200, height = 550 )

manhattan(x = merge_continous_Ewas, chr = "CHR", bp = "MAPINFO", p = "P.value", genomewideline = -log10(2.147985e-05),suggestiveline = FALSE, col = c("blue", "red"))

dev.off()

jpeg(filename = "qq_plot_ewas_binary.jpg")

qq(merge_binary_Ewas$P.value)

dev.off()

jpeg(filename = "qq_plot_ewas_continous.jpg")

qq(merge_continous_Ewas$P.value)

dev.off()

#### Variation Partition analyses:

setwd("C:/GEM_THESE/Final_distangle_EWAS_GWAS/Varpart")

library(vegan)

\#

\# ###load the genotype file with no NA

\# G_df2 <- read.delim("C:/GEM_THESE/Final_distangle_EWAS_GWAS/Final_Variation_partiton/genotype_220_varpart_input_no_NA.txt")

\#

\#

\#

\# #### load the saved file

\# df_E <- read.delim("C:/GEM_THESE/Final_distangle_EWAS_GWAS/epigenotype_220_varpart_input_no_NA.txt")

\#

\# #### prepare the file for the PCA

\# data <- df_E

\# rnames <- data[[1]]# assign labels in column 1 to "rnames"

\# mat_data <- data.matrix(data[,2:221]) # transform column 2 - end into a matrix

\# rownames(mat_data) <- rnames

\# data <- as.matrix(mat_data)

\# data2 <- data/(100)

\# EE <- data2

\#

\# ###transpnse the dataframe

\# EEE <- t(EE)

\# m_E <- data.frame (EEE)

\#

\#

\# df2_G <- G_df2[, names(df_E)]

\#

\# data <- df2_G

\# rnames <- data[[1]]# assign labels in column 1 to "rnames"

\# mat_data <- data.matrix(data[,2:221]) # transform column 2 - end into a matrix

\# rownames(mat_data) <- rnames

\# data <- as.matrix(mat_data)

\# GG <- data

\# GGG <- t(GG)

\# m_G_1 <- data.frame (GGG)

\### make the PCA

\# meth.pc=prcomp(m_E)

\# save(meth.pc, file="meth.pc.Rdata")

load("meth.pc.Rdata")

summary(meth.pc)

\# meth.bs=meth.pc$x[,1:220]

\# write.table(meth.bs, file ="prcomp_epigenotype_220_varpart_input.txt", sep = "\t",row.names = T)

\#

meth.bs <- read.delim("C:/GEM_THESE/Final_distangle_EWAS_GWAS/Varpart/prcomp_epigenotype_220_varpart_input.txt")

\#

\#

\# genet.pc=prcomp(m_G_1)

\# save(genet.pc, file="genet.pc.Rdata")

load("genet.pc.Rdata")

summary(genet.pc)

\# genet.bs <- genet.pc$x[,1:220]

\#

\# write.table(genet.bs, file ="prcomp_genotype_220_varpart_input.txt", sep = "\t", row.names = T)

genet.bs <- read.delim("C:/GEM_THESE/Final_distangle_EWAS_GWAS/Varpart/prcomp_genotype_220_varpart_input.txt")

\# Final_phenotype_binary_220 <- read.delim("C:/GEM_THESE/Final_distangle_EWAS_GWAS/Varpart/Final_phenotype_binary_220.txt")

\#

\# df_PP <- Final_phenotype_binary_220[, names(df_E)]

\#

\#

\# data <- df_PP

\# rnames <- data[[1]]# assign labels in column 1 to "rnames"

\# mat_data <- data.matrix(data[,2:221]) # transform column 2 - end into a matrix

\# rownames(mat_data) <- rnames

\# data <- as.matrix(mat_data)

\# PP <- data

\# PPP <- t(PP)

\# m_P <- data.frame (PPP)

\#

\# write.table(m_P, file ="phenotype_bin_220_varpart_input.txt", sep = "\t", row.names = T)

m_P <- read.delim("C:/GEM_THESE/Final_distangle_EWAS_GWAS/Varpart/phenotype_bin_220_varpart_input.txt")

\# Final_phenotype_continous_220 <- read.delim("C:/GEM_THESE/Final_distangle_EWAS_GWAS/Varpart/Final_phenotype_continous_220.txt")

\# df_PP_con <- Final_phenotype_continous_220[, names(df_E)]

\#

\#

\#

\# data <- df_PP_con

\# rnames <- data[[1]]# assign labels in column 1 to "rnames"

\# mat_data <- data.matrix(data[,2:221]) # transform column 2 - end into a matrix

\# rownames(mat_data) <- rnames

\# data <- as.matrix(mat_data)

\# PP <- data

\# PPP <- t(PP)

\# m_P_con <- data.frame (PPP)

\#

\# write.table(m_P_con, file ="phenotype_con_220_varpart_input.txt", sep = "\t", row.names = T)

m_P_con <- read.delim("C:/GEM_THESE/Final_distangle_EWAS_GWAS/Varpart/phenotype_con_220_varpart_input.txt")

mod0=rda(m_P~1)

mod1=rda(m_P~genet.bs[, 1]+genet.bs[, 2]+genet.bs[, 3]+genet.bs[, 4]+genet.bs[, 5]+genet.bs[, 6]+genet.bs[, 7]+genet.bs[, 8]+genet.bs[, 9]+genet.bs[, 10]+genet.bs[, 11]+genet.bs[, 12]+genet.bs[, 13]+genet.bs[, 14]+genet.bs[, 15]+genet.bs[, 16]+genet.bs[, 17]+genet.bs[, 18]+genet.bs[, 19]+genet.bs[, 20]+genet.bs[, 21]+genet.bs[, 22]+genet.bs[, 23]+genet.bs[, 24]+genet.bs[, 25]+genet.bs[, 26]+genet.bs[, 27]+genet.bs[, 28]+genet.bs[, 29]+genet.bs[, 30]+genet.bs[, 31]+genet.bs[, 32]+genet.bs[, 33]+genet.bs[, 34]+genet.bs[, 35]+genet.bs[, 36]+genet.bs[, 37]+genet.bs[, 38]+genet.bs[, 39]+genet.bs[, 40]+genet.bs[, 41]+genet.bs[, 42]+genet.bs[, 43]+genet.bs[, 44]+genet.bs[, 45]+genet.bs[, 46]+genet.bs[, 47]+genet.bs[, 48]+genet.bs[, 49]+genet.bs[, 50]+genet.bs[, 51]+genet.bs[, 52]+genet.bs[, 53]+genet.bs[, 54]+genet.bs[, 55]+genet.bs[, 56]+genet.bs[, 57]+genet.bs[, 58]+genet.bs[, 59]+genet.bs[, 60]+genet.bs[, 61]+genet.bs[, 62]+genet.bs[, 63]+genet.bs[, 64]+genet.bs[, 65]+genet.bs[, 66]+genet.bs[, 67]+genet.bs[, 68]+genet.bs[, 69]+genet.bs[, 70]+genet.bs[, 71]+genet.bs[, 72]+genet.bs[, 73]+genet.bs[, 74]+genet.bs[, 75]+genet.bs[, 76]+genet.bs[, 77]+genet.bs[, 78]+genet.bs[, 79]+genet.bs[, 80]+genet.bs[, 81]+genet.bs[, 82]+genet.bs[, 83]+genet.bs[, 84]+genet.bs[, 85]+genet.bs[, 86]+genet.bs[, 87]+genet.bs[, 88]+genet.bs[, 89]+genet.bs[, 90]+genet.bs[, 91]+genet.bs[, 92]+genet.bs[, 93]+genet.bs[, 94]+genet.bs[, 95]+genet.bs[, 96]+genet.bs[, 97]+genet.bs[, 98]+genet.bs[, 99]+genet.bs[, 100]+genet.bs[, 101]+genet.bs[, 102]+genet.bs[, 103]+genet.bs[, 104]+genet.bs[, 105]+genet.bs[, 106]+genet.bs[, 107]+genet.bs[, 108]+genet.bs[, 109]+genet.bs[, 110]+genet.bs[, 111]+genet.bs[, 112]+genet.bs[, 113]+genet.bs[, 114]+genet.bs[, 115]+genet.bs[, 116]+genet.bs[, 117]+genet.bs[, 118]+genet.bs[, 119]+genet.bs[, 120]+genet.bs[, 121]+genet.bs[, 122]+genet.bs[, 123]+genet.bs[, 124]+genet.bs[, 125]+genet.bs[, 126]+genet.bs[, 127]+genet.bs[, 128]+genet.bs[, 129]+genet.bs[, 130]+genet.bs[, 131]+genet.bs[, 132]+genet.bs[, 133]+genet.bs[, 134]+genet.bs[, 135]+genet.bs[, 136]+genet.bs[, 137]+genet.bs[, 138]+genet.bs[, 139]+genet.bs[, 140]+genet.bs[, 141]+genet.bs[, 142]+genet.bs[, 143]+genet.bs[, 144]+genet.bs[, 145]+genet.bs[, 146]+genet.bs[, 147]+genet.bs[, 148]+genet.bs[, 149]+genet.bs[, 150]+genet.bs[, 151]+genet.bs[, 152]+genet.bs[, 153]+genet.bs[, 154]+genet.bs[, 155]+genet.bs[, 156]+genet.bs[, 157]+genet.bs[, 158]+genet.bs[, 159]+genet.bs[, 160]+genet.bs[, 161]+genet.bs[, 162]+genet.bs[, 163]+genet.bs[, 164]+genet.bs[, 165]+genet.bs[, 166]+genet.bs[, 167]+genet.bs[, 168]+genet.bs[, 169]+genet.bs[, 170]+genet.bs[, 171]+genet.bs[, 172]+genet.bs[, 173]+genet.bs[, 174]+genet.bs[, 175]+genet.bs[, 176]+genet.bs[, 177]+genet.bs[, 178]+genet.bs[, 179]+genet.bs[, 180]+genet.bs[, 181]+genet.bs[, 182]+genet.bs[, 183]+genet.bs[, 184]+genet.bs[, 185]+genet.bs[, 186]+genet.bs[, 187]+genet.bs[, 188]+genet.bs[, 189]+genet.bs[, 190]+genet.bs[, 191]+genet.bs[, 192]+genet.bs[, 193]+genet.bs[, 194]+genet.bs[, 195]+genet.bs[, 196]+genet.bs[, 197]+genet.bs[, 198]+genet.bs[, 199]+genet.bs[, 200]+genet.bs[, 201]+genet.bs[, 202]+genet.bs[, 203]+genet.bs[, 204]+genet.bs[, 205]+genet.bs[, 206]+genet.bs[, 207]+genet.bs[, 208]+genet.bs[, 209]+genet.bs[, 210]+genet.bs[, 211]+genet.bs[, 212]+genet.bs[, 213]+genet.bs[, 214]+genet.bs[, 215]+genet.bs[, 216]+genet.bs[, 217]+genet.bs[, 218])

###ordistep for binary with all the PCs, here almost most of PCs are significant

gen_bin= ordistep(mod0, mod1, Pin=0.05, permutations=999)

save(gen_bin, file="ordistep_gen_bin.Rdata")

load("ordistep_gen_bin.Rdata")

\### WITH 999 PERM, select the significant axis that been selected by ordistep

GENET=data.frame(cbind(genet.bs[, 151] ,genet.bs[, 126] ,genet.bs[, 28] ,genet.bs[, 42] ,genet.bs[, 178] ,genet.bs[, 35] ,genet.bs[, 102] ,genet.bs[, 166] ,genet.bs[, 149] ,genet.bs[, 209] ,genet.bs[, 135] ,genet.bs[, 95] ,genet.bs[, 6] ,genet.bs[, 156] ,genet.bs[, 73] ,genet.bs[, 158] ,genet.bs[, 23] ,genet.bs[, 133] ,genet.bs[, 21] ,genet.bs[, 199] ,genet.bs[, 183] ,genet.bs[, 39] ,genet.bs[, 5] ,genet.bs[, 116] ,genet.bs[, 177] ,genet.bs[, 186] ,genet.bs[, 86] ,genet.bs[, 88] ,genet.bs[, 90] ,genet.bs[, 34]))

\#

mod2=rda(m_P~meth.bs[, 1]+meth.bs[, 2]+meth.bs[, 3]+meth.bs[, 4]+meth.bs[, 5]+meth.bs[, 6]+meth.bs[, 7]+meth.bs[, 8]+meth.bs[, 9]+meth.bs[, 10]+meth.bs[, 11]+meth.bs[, 12]+meth.bs[, 13]+meth.bs[, 14]+meth.bs[, 15]+meth.bs[, 16]+meth.bs[, 17]+meth.bs[, 18]+meth.bs[, 19]+meth.bs[, 20]+meth.bs[, 21]+meth.bs[, 22]+meth.bs[, 23]+meth.bs[, 24]+meth.bs[, 25]+meth.bs[, 26]+meth.bs[, 27]+meth.bs[, 28]+meth.bs[, 29]+meth.bs[, 30]+meth.bs[, 31]+meth.bs[, 32]+meth.bs[, 33]+meth.bs[, 34]+meth.bs[, 35]+meth.bs[, 36]+meth.bs[, 37]+meth.bs[, 38]+meth.bs[, 39]+meth.bs[, 40]+meth.bs[, 41]+meth.bs[, 42]+meth.bs[, 43]+meth.bs[, 44]+meth.bs[, 45]+meth.bs[, 46]+meth.bs[, 47]+meth.bs[, 48]+meth.bs[, 49]+meth.bs[, 50]+meth.bs[, 51]+meth.bs[, 52]+meth.bs[, 53]+meth.bs[, 54]+meth.bs[, 55]+meth.bs[, 56]+meth.bs[, 57]+meth.bs[, 58]+meth.bs[, 59]+meth.bs[, 60]+meth.bs[, 61]+meth.bs[, 62]+meth.bs[, 63]+meth.bs[, 64]+meth.bs[, 65]+meth.bs[, 66]+meth.bs[, 67]+meth.bs[, 68]+meth.bs[, 69]+meth.bs[, 70]+meth.bs[, 71]+meth.bs[, 72]+meth.bs[, 73]+meth.bs[, 74]+meth.bs[, 75]+meth.bs[, 76]+meth.bs[, 77]+meth.bs[, 78]+meth.bs[, 79]+meth.bs[, 80]+meth.bs[, 81]+meth.bs[, 82]+meth.bs[, 83]+meth.bs[, 84]+meth.bs[, 85]+meth.bs[, 86]+meth.bs[, 87]+meth.bs[, 88]+meth.bs[, 89]+meth.bs[, 90]+meth.bs[, 91]+meth.bs[, 92]+meth.bs[, 93]+meth.bs[, 94]+meth.bs[, 95]+meth.bs[, 96]+meth.bs[, 97]+meth.bs[, 98]+meth.bs[, 99]+meth.bs[, 100]+meth.bs[, 101]+meth.bs[, 102]+meth.bs[, 103]+meth.bs[, 104]+meth.bs[, 105]+meth.bs[, 106]+meth.bs[, 107]+meth.bs[, 108]+meth.bs[, 109]+meth.bs[, 110]+meth.bs[, 111]+meth.bs[, 112]+meth.bs[, 113]+meth.bs[, 114]+meth.bs[, 115]+meth.bs[, 116]+meth.bs[, 117]+meth.bs[, 118]+meth.bs[, 119]+meth.bs[, 120]+meth.bs[, 121]+meth.bs[, 122]+meth.bs[, 123]+meth.bs[, 124]+meth.bs[, 125]+meth.bs[, 126]+meth.bs[, 127]+meth.bs[, 128]+meth.bs[, 129]+meth.bs[, 130]+meth.bs[, 131]+meth.bs[, 132]+meth.bs[, 133]+meth.bs[, 134]+meth.bs[, 135]+meth.bs[, 136]+meth.bs[, 137]+meth.bs[, 138]+meth.bs[, 139]+meth.bs[, 140]+meth.bs[, 141]+meth.bs[, 142]+meth.bs[, 143]+meth.bs[, 144]+meth.bs[, 145]+meth.bs[, 146]+meth.bs[, 147]+meth.bs[, 148]+meth.bs[, 149]+meth.bs[, 150]+meth.bs[, 151]+meth.bs[, 152]+meth.bs[, 153]+meth.bs[, 154]+meth.bs[, 155]+meth.bs[, 156]+meth.bs[, 157]+meth.bs[, 158]+meth.bs[, 159]+meth.bs[, 160]+meth.bs[, 161]+meth.bs[, 162]+meth.bs[, 163]+meth.bs[, 164]+meth.bs[, 165]+meth.bs[, 166]+meth.bs[, 167]+meth.bs[, 168]+meth.bs[, 169]+meth.bs[, 170]+meth.bs[, 171]+meth.bs[, 172]+meth.bs[, 173]+meth.bs[, 174]+meth.bs[, 175]+meth.bs[, 176]+meth.bs[, 177]+meth.bs[, 178]+meth.bs[, 179]+meth.bs[, 180]+meth.bs[, 181]+meth.bs[, 182]+meth.bs[, 183]+meth.bs[, 184]+meth.bs[, 185]+meth.bs[, 186]+meth.bs[, 187]+meth.bs[, 188]+meth.bs[, 189]+meth.bs[, 190]+meth.bs[, 191]+meth.bs[, 192]+meth.bs[, 193]+meth.bs[, 194]+meth.bs[, 195]+meth.bs[, 196]+meth.bs[, 197]+meth.bs[, 198]+meth.bs[, 199]+meth.bs[, 200]+meth.bs[, 201]+meth.bs[, 202]+meth.bs[, 203]+meth.bs[, 204]+meth.bs[, 205]+meth.bs[, 206]+meth.bs[, 207]+meth.bs[, 208]+meth.bs[, 209]+meth.bs[, 210]+meth.bs[, 211]+meth.bs[, 212]+meth.bs[, 213]+meth.bs[, 214]+meth.bs[, 215]+meth.bs[, 216]+meth.bs[, 217]+meth.bs[, 218])

anova(mod2)

\#

meth_bin= ordistep(mod0, mod2, Pin=0.05, permutations=999)

save(meth_bin, file="ordistep_meth_bin.Rdata")

load("ordistep_meth_bin.Rdata")

\### WITH 999 PERM, select the significant axis that been selected by ordistep

METH=data.frame(cbind(meth.bs[, 2] ,meth.bs[, 6] ,meth.bs[, 4] ,meth.bs[, 35] ,meth.bs[, 24] ,meth.bs[, 1] ,meth.bs[, 72] ,meth.bs[, 15] ,meth.bs[, 100] ,meth.bs[, 106] ,meth.bs[, 86] ,meth.bs[, 26] ,meth.bs[, 77] ,meth.bs[, 22] ,meth.bs[, 145] ,meth.bs[, 208] ,meth.bs[, 141] ,meth.bs[, 200] ,meth.bs[, 5] ,meth.bs[, 43] ,meth.bs[, 32] ,meth.bs[, 13] ,meth.bs[, 158] ,meth.bs[, 197] ,meth.bs[, 81] ,meth.bs[, 108] ,meth.bs[, 153] ,meth.bs[, 20] ,meth.bs[, 121] ,meth.bs[, 148] ,meth.bs[, 55] ,meth.bs[, 149] ,meth.bs[, 103] ,meth.bs[, 90] ,meth.bs[, 94] ,meth.bs[, 129] ,meth.bs[, 82] ,meth.bs[, 83] ,meth.bs[, 68] ,meth.bs[, 87]))

varpart_bin = varpart(m_P,GENET,METH)

varpart_bin

\# Partition of variance in RDA

\#

\# Call: varpart(Y = m_P, X = GENET, METH)

\#

\# Explanatory tables:

\#	X1: GENET

\# X2: METH

\#

\# No. of explanatory tables: 2

\# Total variation (SS): 54.709

\# Variance: 0.24981

\# No. of observations: 220

\#

\# Partition table:

\#	Df R.squared Adj.R.squared Testable

\# [a+b] = X1	30	0.53922	0.46608	TRUE

\# [b+c] = X2	40	0.66981	0.59602	TRUE

\# [a+b+c] = X1+X2	70	0.81431	0.72707	TRUE

\# Individual fractions

\# [a] = X1|X2	30	0.13105	TRUE

\# [b]	0	0.33503	FALSE

\# [c] = X2|X1	40	0.26098	TRUE

\# [d] = Residuals	0.27293	FALSE

\# ---

\#	Use function 'rda' to test significance of fractions of interest

plot(varpart_bin, digits = 1, Xnames = c('Genetic', 'Epigenetic'), bg = c('Blue', 'red'))

################## With semi-continuous phenotype

\#

modA=rda(m_P_con~1)

modB=rda(m_P_con~genet.bs[, 1]+genet.bs[, 2]+genet.bs[, 3]+genet.bs[, 4]+genet.bs[, 5]+genet.bs[, 6]+genet.bs[, 7]+genet.bs[, 8]+genet.bs[, 9]+genet.bs[, 10]+genet.bs[, 11]+genet.bs[, 12]+genet.bs[, 13]+genet.bs[, 14]+genet.bs[, 15]+genet.bs[, 16]+genet.bs[, 17]+genet.bs[, 18]+genet.bs[, 19]+genet.bs[, 20]+genet.bs[, 21]+genet.bs[, 22]+genet.bs[, 23]+genet.bs[, 24]+genet.bs[, 25]+genet.bs[, 26]+genet.bs[, 27]+genet.bs[, 28]+genet.bs[, 29]+genet.bs[, 30]+genet.bs[, 31]+genet.bs[, 32]+genet.bs[, 33]+genet.bs[, 34]+genet.bs[, 35]+genet.bs[, 36]+genet.bs[, 37]+genet.bs[, 38]+genet.bs[, 39]+genet.bs[, 40]+genet.bs[, 41]+genet.bs[, 42]+genet.bs[, 43]+genet.bs[, 44]+genet.bs[, 45]+genet.bs[, 46]+genet.bs[, 47]+genet.bs[, 48]+genet.bs[, 49]+genet.bs[, 50]+genet.bs[, 51]+genet.bs[, 52]+genet.bs[, 53]+genet.bs[, 54]+genet.bs[, 55]+genet.bs[, 56]+genet.bs[, 57]+genet.bs[, 58]+genet.bs[, 59]+genet.bs[, 60]+genet.bs[, 61]+genet.bs[, 62]+genet.bs[, 63]+genet.bs[, 64]+genet.bs[, 65]+genet.bs[, 66]+genet.bs[, 67]+genet.bs[, 68]+genet.bs[, 69]+genet.bs[, 70]+genet.bs[, 71]+genet.bs[, 72]+genet.bs[, 73]+genet.bs[, 74]+genet.bs[, 75]+genet.bs[, 76]+genet.bs[, 77]+genet.bs[, 78]+genet.bs[, 79]+genet.bs[, 80]+genet.bs[, 81]+genet.bs[, 82]+genet.bs[, 83]+genet.bs[, 84]+genet.bs[, 85]+genet.bs[, 86]+genet.bs[, 87]+genet.bs[, 88]+genet.bs[, 89]+genet.bs[, 90]+genet.bs[, 91]+genet.bs[, 92]+genet.bs[, 93]+genet.bs[, 94]+genet.bs[, 95]+genet.bs[, 96]+genet.bs[, 97]+genet.bs[, 98]+genet.bs[, 99]+genet.bs[, 100]+genet.bs[, 101]+genet.bs[, 102]+genet.bs[, 103]+genet.bs[, 104]+genet.bs[, 105]+genet.bs[, 106]+genet.bs[, 107]+genet.bs[, 108]+genet.bs[, 109]+genet.bs[, 110]+genet.bs[, 111]+genet.bs[, 112]+genet.bs[, 113]+genet.bs[, 114]+genet.bs[, 115]+genet.bs[, 116]+genet.bs[, 117]+genet.bs[, 118]+genet.bs[, 119]+genet.bs[, 120]+genet.bs[, 121]+genet.bs[, 122]+genet.bs[, 123]+genet.bs[, 124]+genet.bs[, 125]+genet.bs[, 126]+genet.bs[, 127]+genet.bs[, 128]+genet.bs[, 129]+genet.bs[, 130]+genet.bs[, 131]+genet.bs[, 132]+genet.bs[, 133]+genet.bs[, 134]+genet.bs[, 135]+genet.bs[, 136]+genet.bs[, 137]+genet.bs[, 138]+genet.bs[, 139]+genet.bs[, 140]+genet.bs[, 141]+genet.bs[, 142]+genet.bs[, 143]+genet.bs[, 144]+genet.bs[, 145]+genet.bs[, 146]+genet.bs[, 147]+genet.bs[, 148]+genet.bs[, 149]+genet.bs[, 150]+genet.bs[, 151]+genet.bs[, 152]+genet.bs[, 153]+genet.bs[, 154]+genet.bs[, 155]+genet.bs[, 156]+genet.bs[, 157]+genet.bs[, 158]+genet.bs[, 159]+genet.bs[, 160]+genet.bs[, 161]+genet.bs[, 162]+genet.bs[, 163]+genet.bs[, 164]+genet.bs[, 165]+genet.bs[, 166]+genet.bs[, 167]+genet.bs[, 168]+genet.bs[, 169]+genet.bs[, 170]+genet.bs[, 171]+genet.bs[, 172]+genet.bs[, 173]+genet.bs[, 174]+genet.bs[, 175]+genet.bs[, 176]+genet.bs[, 177]+genet.bs[, 178]+genet.bs[, 179]+genet.bs[, 180]+genet.bs[, 181]+genet.bs[, 182]+genet.bs[, 183]+genet.bs[, 184]+genet.bs[, 185]+genet.bs[, 186]+genet.bs[, 187]+genet.bs[, 188]+genet.bs[, 189]+genet.bs[, 190]+genet.bs[, 191]+genet.bs[, 192]+genet.bs[, 193]+genet.bs[, 194]+genet.bs[, 195]+genet.bs[, 196]+genet.bs[, 197]+genet.bs[, 198]+genet.bs[, 199]+genet.bs[, 200]+genet.bs[, 201]+genet.bs[, 202]+genet.bs[, 203]+genet.bs[, 204]+genet.bs[, 205]+genet.bs[, 206]+genet.bs[, 207]+genet.bs[, 208]+genet.bs[, 209]+genet.bs[, 210]+genet.bs[, 211]+genet.bs[, 212]+genet.bs[, 213]+genet.bs[, 214]+genet.bs[, 215]+genet.bs[, 216]+genet.bs[, 217]+genet.bs[, 218])

\#

\#

###ordistep for binary with all the PCs, here almost most of PCs are significant

genet_con = ordistep(modA, modB, Pin=0.05, permutations=999)

save(genet_con, file="ordistep_genet_con.Rdata")

genet_con$anova

\### WITH 999 PERM, select the significant axis that been selected by ordistep

GENET2=data.frame(cbind(genet.bs[, 35] ,genet.bs[, 151] ,genet.bs[, 28] ,genet.bs[, 126] ,genet.bs[, 42] ,genet.bs[, 73] ,genet.bs[, 21] ,genet.bs[, 178] ,genet.bs[, 102] ,genet.bs[, 158] ,genet.bs[, 166] ,genet.bs[, 23] ,genet.bs[, 45] ,genet.bs[, 6] ,genet.bs[, 209] ,genet.bs[, 146] ,genet.bs[, 55] ,genet.bs[, 5] ,genet.bs[, 133] ,genet.bs[, 116] ,genet.bs[, 95] ,genet.bs[, 135] ,genet.bs[, 118] ,genet.bs[, 149] ,genet.bs[, 215] ,genet.bs[, 205] ,genet.bs[, 72] ,genet.bs[, 159] ,genet.bs[, 98]))

\#

modC=rda(m_P_con~meth.bs[, 1]+meth.bs[, 2]+meth.bs[, 3]+meth.bs[, 4]+meth.bs[, 5]+meth.bs[, 6]+meth.bs[, 7]+meth.bs[, 8]+meth.bs[, 9]+meth.bs[, 10]+meth.bs[, 11]+meth.bs[, 12]+meth.bs[, 13]+meth.bs[, 14]+meth.bs[, 15]+meth.bs[, 16]+meth.bs[, 17]+meth.bs[, 18]+meth.bs[, 19]+meth.bs[, 20]+meth.bs[, 21]+meth.bs[, 22]+meth.bs[, 23]+meth.bs[, 24]+meth.bs[, 25]+meth.bs[, 26]+meth.bs[, 27]+meth.bs[, 28]+meth.bs[, 29]+meth.bs[, 30]+meth.bs[, 31]+meth.bs[, 32]+meth.bs[, 33]+meth.bs[, 34]+meth.bs[, 35]+meth.bs[, 36]+meth.bs[, 37]+meth.bs[, 38]+meth.bs[, 39]+meth.bs[, 40]+meth.bs[, 41]+meth.bs[, 42]+meth.bs[, 43]+meth.bs[, 44]+meth.bs[, 45]+meth.bs[, 46]+meth.bs[, 47]+meth.bs[, 48]+meth.bs[, 49]+meth.bs[, 50]+meth.bs[, 51]+meth.bs[, 52]+meth.bs[, 53]+meth.bs[, 54]+meth.bs[, 55]+meth.bs[, 56]+meth.bs[, 57]+meth.bs[, 58]+meth.bs[, 59]+meth.bs[, 60]+meth.bs[, 61]+meth.bs[, 62]+meth.bs[, 63]+meth.bs[, 64]+meth.bs[, 65]+meth.bs[, 66]+meth.bs[, 67]+meth.bs[, 68]+meth.bs[, 69]+meth.bs[, 70]+meth.bs[, 71]+meth.bs[, 72]+meth.bs[, 73]+meth.bs[, 74]+meth.bs[, 75]+meth.bs[, 76]+meth.bs[, 77]+meth.bs[, 78]+meth.bs[, 79]+meth.bs[, 80]+meth.bs[, 81]+meth.bs[, 82]+meth.bs[, 83]+meth.bs[, 84]+meth.bs[, 85]+meth.bs[, 86]+meth.bs[, 87]+meth.bs[, 88]+meth.bs[, 89]+meth.bs[, 90]+meth.bs[, 91]+meth.bs[, 92]+meth.bs[, 93]+meth.bs[, 94]+meth.bs[, 95]+meth.bs[, 96]+meth.bs[, 97]+meth.bs[, 98]+meth.bs[, 99]+meth.bs[, 100]+meth.bs[, 101]+meth.bs[, 102]+meth.bs[, 103]+meth.bs[, 104]+meth.bs[, 105]+meth.bs[, 106]+meth.bs[, 107]+meth.bs[, 108]+meth.bs[, 109]+meth.bs[, 110]+meth.bs[, 111]+meth.bs[, 112]+meth.bs[, 113]+meth.bs[, 114]+meth.bs[, 115]+meth.bs[, 116]+meth.bs[, 117]+meth.bs[, 118]+meth.bs[, 119]+meth.bs[, 120]+meth.bs[, 121]+meth.bs[, 122]+meth.bs[, 123]+meth.bs[, 124]+meth.bs[, 125]+meth.bs[, 126]+meth.bs[, 127]+meth.bs[, 128]+meth.bs[, 129]+meth.bs[, 130]+meth.bs[, 131]+meth.bs[, 132]+meth.bs[, 133]+meth.bs[, 134]+meth.bs[, 135]+meth.bs[, 136]+meth.bs[, 137]+meth.bs[, 138]+meth.bs[, 139]+meth.bs[, 140]+meth.bs[, 141]+meth.bs[, 142]+meth.bs[, 143]+meth.bs[, 144]+meth.bs[, 145]+meth.bs[, 146]+meth.bs[, 147]+meth.bs[, 148]+meth.bs[, 149]+meth.bs[, 150]+meth.bs[, 151]+meth.bs[, 152]+meth.bs[, 153]+meth.bs[, 154]+meth.bs[, 155]+meth.bs[, 156]+meth.bs[, 157]+meth.bs[, 158]+meth.bs[, 159]+meth.bs[, 160]+meth.bs[, 161]+meth.bs[, 162]+meth.bs[, 163]+meth.bs[, 164]+meth.bs[, 165]+meth.bs[, 166]+meth.bs[, 167]+meth.bs[, 168]+meth.bs[, 169]+meth.bs[, 170]+meth.bs[, 171]+meth.bs[, 172]+meth.bs[, 173]+meth.bs[, 174]+meth.bs[, 175]+meth.bs[, 176]+meth.bs[, 177]+meth.bs[, 178]+meth.bs[, 179]+meth.bs[, 180]+meth.bs[, 181]+meth.bs[, 182]+meth.bs[, 183]+meth.bs[, 184]+meth.bs[, 185]+meth.bs[, 186]+meth.bs[, 187]+meth.bs[, 188]+meth.bs[, 189]+meth.bs[, 190]+meth.bs[, 191]+meth.bs[, 192]+meth.bs[, 193]+meth.bs[, 194]+meth.bs[, 195]+meth.bs[, 196]+meth.bs[, 197]+meth.bs[, 198]+meth.bs[, 199]+meth.bs[, 200]+meth.bs[, 201]+meth.bs[, 202]+meth.bs[, 203]+meth.bs[, 204]+meth.bs[, 205]+meth.bs[, 206]+meth.bs[, 207]+meth.bs[, 208]+meth.bs[, 209]+meth.bs[, 210]+meth.bs[, 211]+meth.bs[, 212]+meth.bs[, 213]+meth.bs[, 214]+meth.bs[, 215]+meth.bs[, 216]+meth.bs[, 217]+meth.bs[, 218])

\#

\#

\#

meth_con = ordistep(modA, modC, Pin=0.05, permutations=999)

save(meth_con, file="ordistep_meth_con.Rdata")

meth_con$anova

\### WITH 999 PERM, select the significant axis that been selected by ordistep

METH2=data.frame(cbind(meth.bs[, 4] ,meth.bs[, 2] ,meth.bs[, 1] ,meth.bs[, 6] ,meth.bs[, 24] ,meth.bs[, 35] ,meth.bs[, 15] ,meth.bs[, 5] ,meth.bs[, 100] ,meth.bs[, 106] ,meth.bs[, 22] ,meth.bs[, 86] ,meth.bs[, 145] ,meth.bs[, 32] ,meth.bs[, 72] ,meth.bs[, 141] ,meth.bs[, 153] ,meth.bs[, 26] ,meth.bs[, 108] ,meth.bs[, 177] ,meth.bs[, 30] ,meth.bs[, 139] ,meth.bs[, 66] ,meth.bs[, 202] ,meth.bs[, 103] ,meth.bs[, 77] ,meth.bs[, 112] ,meth.bs[, 82] ,meth.bs[, 119]))

varpart_con = varpart(m_P_con,GENET2,METH2)

varpart_con

\# Partition of variance in RDA

\#

\# Call: varpart(Y = m_P_con, X = GENET2, METH2)

\#

\# Explanatory tables:

\#	X1: GENET2

\# X2: METH2

\#

\# No. of explanatory tables: 2

\# Total variation (SS): 2627146

\# Variance: 11996

\# No. of observations: 220

\#

\# Partition table:

\#	Df R.squared Adj.R.squared Testable

\# [a+b] = X1	29	0.55111	0.48259	TRUE

\# [b+c] = X2	29	0.57945	0.51527	TRUE

\# [a+b+c] = X1+X2	58	0.74700	0.65586	TRUE

\# Individual fractions

\# [a] = X1|X2	29	0.14059	TRUE

\# [b]	0	0.34200	FALSE

\# [c] = X2|X1	29	0.17327	TRUE

\# [d] = Residuals	0.34414	FALSE

\# ---

\#	Use function 'rda' to test significance of fractions of interest

plot(varpart_con, digits = 1, Xnames =
